# Trisomy 21 Perturbs Oligodendrocyte Developmental States and Oligo–Neuron Communication in Human Forebrain Assembloids

**DOI:** 10.64898/2026.10.30.763166

**Authors:** Natalie Baker Campbell, Sonal Khanna, Michelle J Kim, Elizabeth K Kharitonova, Maya P Weidman, Cody Headings, Nathan X Chai, Clara Chung, Samantha Chung, Grace Field, Miriam Linetsky, Martin Thunemann, Joshua D Campbell, Maria Medalla, Ella Zeldich

## Abstract

Intellectual disability is a hallmark of Down syndrome (DS), caused by the triplication of human chromosome 21. Abnormal neuronal connectivity and impaired myelination are consistently observed in DS, yet the underlying mechanisms remain poorly understood. Here, we used DS-derived isogenic induced pluripotent stem cell-derived forebrain assembloids (FAs) to investigate oligodendrocyte lineage cell (OLLC) development and neuron–glia interactions. Trisomic FAs exhibited an expansion of BCAS1+ pre-myelinating OLLCs, without a corresponding increase in myelin basic protein (MBP) expression. Transcriptomic analyses identified a euploid-enriched, trophic and maturation-supportive state (OLLC_2-TMS) and a trisomy-enriched synaptic interactions and differentiation state (OLLC_3-SID), lacking coordinated enrichment in myelination-promoting programs. Corresponding transcriptional signatures were identified in OLLCs from early postnatal human cortex. Increased excitatory neuronal contacts with pre-myelinating OLLCs accompanied ultrastructural myelin abnormalities in trisomic assembloids. Our findings suggest that trisomy redirects OLLC development toward a pre-myelinating state, uncoupling oligodendrocyte differentiation from productive myelination.

## Introduction

Down Syndrome (DS) or Trisomy 21 is a genetic disorder caused by the triplication of human chromosome 21 (Hsa21)^1^, resulting in neuroanatomical differences and intellectual disability. Impaired oligodendrocyte development and myelination has been consistently observed in trisomy and is believed to contribute to neuropathology in DS^2–4^. These deficits are reflected in altered expression of maturation markers within oligodendrocyte lineage cells (OLLCs), abnormal organization of myelin fibers, and decreased myelin density in both mouse models of DS and human postmortem brains^2,5,6^. The underlying cause of these deficits, and which developmental stages are affected remains poorly understood, particularly in human cells. It is also unclear whether impaired myelination in DS is primarily results from altered OLLC lineage commitment, a general reduction in OLLC production, disruption of developmental progression, or a specific differentiated state that fails to support productive myelination.

OLLC and neuronal development are closely coupled^7,8^. Oligodendrocyte progenitor cells (OPCs) form synapses with neurons, receive direct synaptic input and respond to neurotransmitter-mediated signaling, including glutamatergic and GABAergic transmission^9–12^. Throughout OLLC development, OLLC–neuronal interactions are modified as oligodendrocytes differentiate and acquire myelinating properties^10,13,14^, including changes in axon-recognition and adhesion, trophic support, and synaptic signaling that supports the establishment of myelinating axon–glia interactions^9,15,16^. Alongside impaired oligodendrocyte development and myelin production, DS disturbs neuronal activity and circuitry from early developmental stages, including alterations in glutamatergic signaling, neuronal excitability and excitatory/inhibitory balance, resulting in decreased dendritic arborization and reduced synaptic connectivity^17–22^. Despite growing recognition that neuron–glia interactions are critical for normal brain development^8,23,24^, the effect of trisomy on oligodendrocyte lineage progression and remodeling of neuron–OLLC interactions remains largely unexplored. Addressing this question in a human genetic context requires a platform in which OLLCs co-develop with neuronal populations and can be examined within a shared physiologically-relevant environment.

OPCs arise in three consecutive developmental waves. The first two waves originate in the ventral telencephalon and migrate dorsally where they intermingle with the third wave that originates postnatally in the dorsal cerebral cortex^25–27^. These distinct developmental origins and migration of OPCs between forebrain regions reflect another essential component of OLLC development required to comprehensively investigate the effect of trisomy on OLLC differentiation and maturation trajectories, as well as the establishment of glia–axonal interactions and myelination.

Here, we generated forebrain assembloids (FAs) by fusing regionally-patterned ventral cortical organoids (vCOs)^28^, and dorsal cortical organoids (dCOs)^29^, containing expanded OLLCs populations. This approach recapitulates OLLC origins, enabling the study of OLLC migration across cortical regions, their progression through differentiation stages, the acquisition of maturation-associated signatures, and their interactions with neuronal populations within a three-dimensional human cellular environment^30–32^. Using phenotypic, transcriptomic, live-cell imaging, and ultrastructural approaches in three isogenic euploid-trisomic paired cell lines, we identified a divergence in OLLC development in trisomic FAs. Our analyses revealed that trisomic OLLCs shift toward a distinct differentiated state associated with enhanced neuronal and synapse-related programs. In contrast, euploid OLLCs preferentially acquire trophic, metabolic, and adhesion-related programs, consistent with a physiological myelination-supportive state. Importantly, this previously unrecognized trisomy-associated differentiated state corresponds to OLLCs identified in transcriptomic data from developing human DS brains. Trisomic FA-derived OLLCs also display preferential synaptic interactions with excitatory neurons and disruptions in myelin ultrastructure. Together, these findings suggest that trisomy shifts oligodendrocyte development away from normal neuron–glia communication toward a state where differentiation programs become uncoupled from programs supporting productive myelination.

## Results

### Trisomy 21 splits OLLC developmental trajectory, expanding the BCAS1+ pre-myelinating OLLC population

To investigate the developmental molecular mechanisms leading to abnormal oligodendrocyte biology and altered myelination in Trisomy 21, we established a human FA model. FAs contain heterogenous cell populations of neuronal and glial cells, including OLLCs across developmental stages. Using three paired isogenic lines, we generated dCOs^29,33,34^ and vCOs^28^, both containing OLLCs, which were subsequently fused into FAs to model oligodendrocyte specification, migration, and maturation in a 3D environment (**Fig. 1a, b**). These FAs recapitulate the sequential, region-specific waves of oligodendrogenesis, in which early ventrally-derived OPCs migrate into and populate the dorsal forebrain, where they are joined by a wave of dorsally-derived OPCs^25,30^.

**Figure 1:**
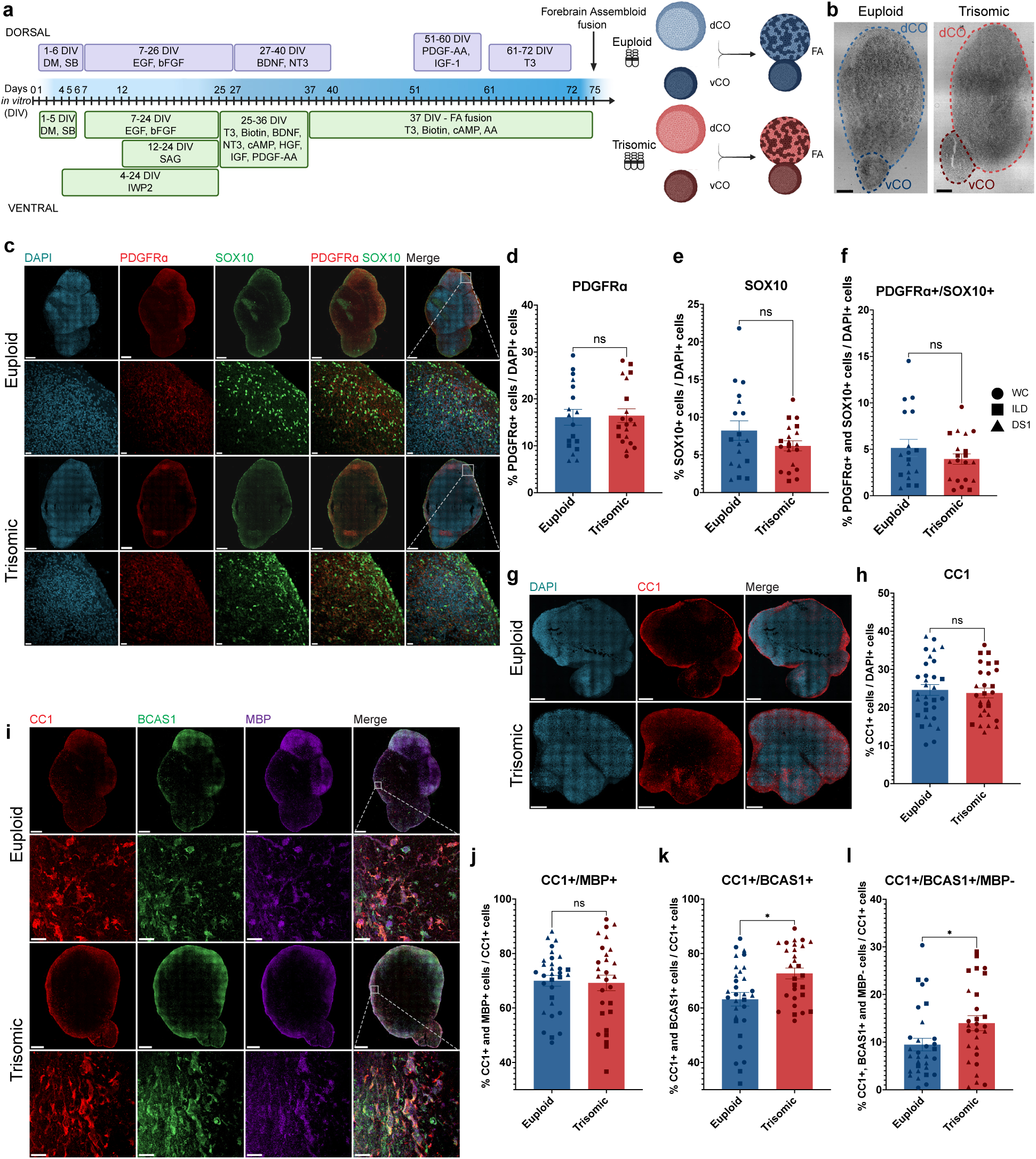
FAs recapitulate regional identities and demonstrate developmental trajectory perturbations in Trisomy 21. **a,** dCO and vCO culturing protocols adapted from Madhavan et al 2018^29^ and Marton et al 2019^28^, and FA fusion schematic adapted from Sloan et al 2018^32^. Created in BioRender. Campbell, NB (2026) https://BioRender.com/lklkxmo **b**, Brightfield images of 150 *DIV* FAs showing vCO and dCO fusion. Scale bar, 200 µm. **c**, Representative stitched and inset confocal images from euploid and trisomic FAs at 150 *DIV* showing PDGFRα (red) and SOX10 (green) expression. Scale bars, 300 µm (overviews) and 20 µm (insets). **d–f,** Quantification of percentage of (**d**) PDGFRα+ cells (LME, β = +0.3454, *p* = 0.8740), (**e**) SOX10+ cells (LME, β = -2.1299, *p* = 0.1125), and (**f**) PDGFRα+/SOX10+ cells (LME, β = -1.16498, *p* = 0.2162) within total DAPI+ cell population per FA. Euploid, *n* = 18. Trisomic, *n* = 20 FAs. **g**, Representative confocal images from euploid and trisomic FAs at 150 *DIV* showing CC1 (red). Scale bar, 300 µm. **h**, Quantification of percentage of CC1+ cells within the total DAPI+ population (LME, β = -0.9875, *p* = 0.6014) per FA. Euploid, *n* = 31. Trisomic, *n* = 28 FAs. **i**, Representative stitched and inset confocal images from euploid and trisomic FAs at 150 *DIV* showing CC1 (red), BCAS1 (green), and MBP (magenta). Scale bars, 300 µm (overviews) and 20 µm (insets). **j–l,** Quantification of percentage of (**j**) CC1+/MBP+ cells (LME, β = +0.2815, *p* = 0.9180), (**k**) CC1+/BCAS1+ cells (LME, β = +10.8061, *p* = 0.0003), and (**l**) CC1+/MBP-/BCAS1+ cells (LME, β = +3.6692, *p* = 0.0459) within total CC1+ cell population per FA. Euploid, *n* = 31. Trisomic, *n* = 28 FAs. All graphs display results from two independent differentiation experiments. All comparisons were analyzed using linear mixed-effects models, with genotype effect reported in the figure legend and the effects of cell line and differentiation summarized in **Supplementary Table 1**. Each point represents one FA and each symbol shape represents an isogenic cell line pair. circle = WC; square = ILD; triangle = DS1. Results shown as mean ± SEM. * *p* < 0.05, ns = not significant.

At 50 days *in vitro* (*DIV)*, prior to FA fusion, dCOs and vCOs expressed regional markers PAX6 and NKX2.1, respectively, confirming dorsal and ventral forebrain patterning in both CO models (**Supplementary Fig. 1a, b**). At this early timepoint, both vCOs and dCOs contained OLLCs expressing the early pan-oligodendrocyte marker, SOX10^35^, with no significant differences in their proportions in dCOs between euploid and trisomic dCOs (*p*=0.8029) or vCOs (*p*=0.4990; **Supplementary Fig. 1c–f**), suggesting trisomy does not result in consistent alterations in early OLLC production, at least as assessed by SOX10 alone, and abnormalities in the oligodendrocyte lineage may arise later in development. All comparisons were analyzed using linear mixed-effects (LME) models, with the effects of cell line and differentiation summarized in **Supplementary Table 1**.

To investigate subsequent developmental events, we examined FAs at 150 *DIV*, the time point at which FAs are expected to contain OLLCs at multiple developmental stages, including OPCs, pre-myelinating and mature oligodendrocytes. Both euploid and trisomic FAs showed robust expression of canonical OPC markers, with no differences in the number of PDGFRα+ (*p*=0.8740), SOX10+ (*p*=0.1125), or PDGFRα+/SOX10+ co-positive cells (*p*=0.2162) between the genotypes (**Fig. 1c–f**). Additionally, the number of cells expressing CC1, a mature oligodendrocyte marker^36–38^, did not differ between the genotypes (*p*=0.6014, **Fig. 1g, h**), suggesting total OLLC production is not affected by trisomy.

We then examined the expression of MBP, necessary for myelin formation^39^, and BCAS1, expressed in transitioning, pre-myelinating oligodendrocytes^40^, within the CC1+ population. We did not detect a difference in the proportion of CC1+ cells co-expressing MBP (CC1+/MBP+) in trisomic FAs (*p*=0.9180; **Fig. 1i, j**). Conversely, trisomic OLLCs demonstrated a significant increase in the pre-myelinating CC1+/BCAS1+ OLLC population (*p*=0.0003; **Fig. 1i, k**). To investigate this population further, we separated CC1+/BCAS1+ cells based on their MBP expression. Within this CC1+/BCAS1+ population, trisomic FAs exhibited a significant increase in the population of CC1+/BCAS1+/MBP-cells (*p*=0.0459; **Fig. 1i, l**,). Together, these results reveal an expansion of the population of trisomic pre-myelinating BCAS1+ OLLCs without a corresponding increase in MBP expression, indicating that trisomy alters the progression of OLLCs along developmental and maturation trajectories.

### Altered OLLC migratory dynamics emerge early in trisomic FAs

The increased population of pre-myelinating CC1+/BCAS1+ OLLCs at 150 *DIV* prompted us to examine whether developmental alterations emerge earlier during OPC migration from ventral to dorsal forebrain regions, and if these alterations affect OLLC distribution. Since OPC migration is an essential part of oligodendrocyte development and deficits in cellular motility have been previously reported in trisomy^41,42^, we examined OLLC migratory dynamics at 120 *DIV* using our recently established live-cell imaging and tracking pipeline^43^. Ventrally-derived OLLCs were virally labeled with *SOX10-tdTomato*^43,44^ prior to FA fusion and their migration into the dorsal compartment was tracked by live-cell imaging over 18 hours (**Fig. 2a–f**).

**Figure 2:**
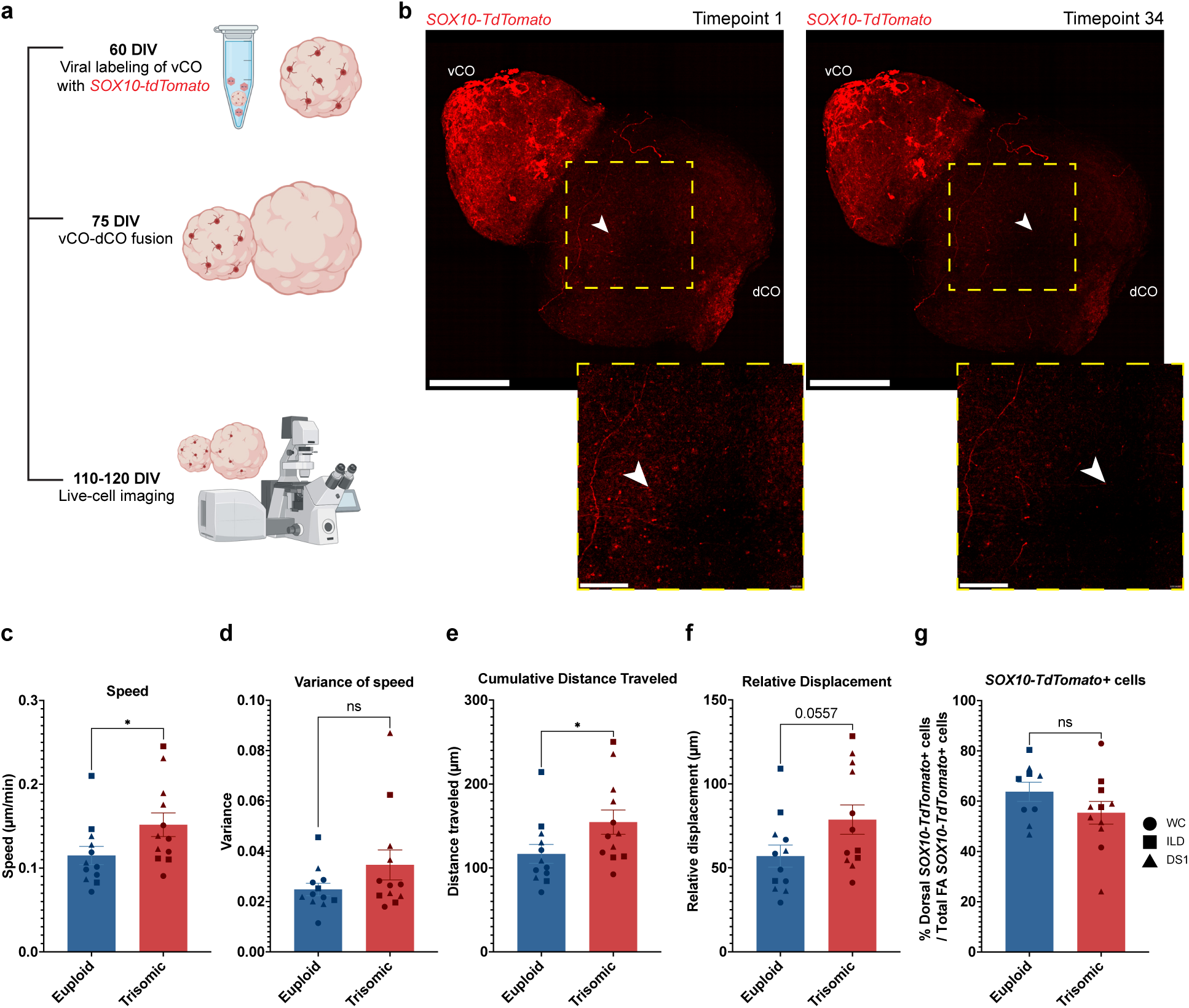
Trisomic OLLCs demonstrate increased speed and distance of migration in human forebrain assembloids. **a**, Experimental design schematic showing viral transduction of vCOs with *SOX10-tdTomato* label, FA fusion, and live-cell imaging. Created in BioRender. Campbell, NB (2026) https://BioRender.com/lklkxmo **b,** Representative high-resolution stitched maximum intensity projection confocal images from live-cell imaging at 120 *DIV* showing FA at first and last timepoint with insets featuring one migrating cell moving throughout the dCO portion. Arrows indicate start and end points of cell movement. Scale bars, 300 µm (overviews) and 50 µm (insets). **c–f,** Quantification of (**c**) speed (µm/min; LME, β = +0.03557, *p* = 0.0430), (**d**) variance of speed (LME, β = +0.0064, *p* = 0.2670), (**e**) cumulative distance (µm; LME, β = +36.5035, *p* = 0.0421), and (**f**) relative displacement (µm; LME, β = +21.0720, *p* = 0.0557) per FA. Euploid, *n* = 12. Trisomic, *n* = 12 FAs. **g**, Quantification of *SOX10-TdTomato*+ cells that have migrated from the vCO to the dCO (LME, β = -1.5095, *p* = 0.8059) within total *SOX10-TdTomato*+ population per FA. Euploid, *n* = 9. Trisomic, *n* = 11 FAs. Graphs display results from two independent differentiation experiments. All comparisons were analyzed using linear mixed-effects models, with genotype effect reported in the figure legend and the effects of cell line and differentiation summarized in **Supplementary Table 1**. Each point represents one FA and each symbol shape represents an isogenic cell line pair. circle = WC; square = ILD; triangle = DS1. Results shown as mean ± SEM. * *p* < 0.05, ns = not significant.

Tracked trisomic *SOX10-tdTomato+* OLLCs displayed significantly increased migration speed (*p=*0.0430; **Fig. 2c**) and cumulative distance traveled (*p=*0.0421; **Fig 2e**), but no difference in the speed variance (*p*=0.2670; **Fig. 2d**), and a trend towards increased relative displacement (*p=*0.0557; **Fig 2f, Supplementary Fig. 2a–d**). Despite these differences in motility, we found no difference in the proportion of vCO-derived *SOX10-tdTomato+* OLLCs in dCO portions when normalized to total *SOX10-tdTomato+* cells per FA (*p*=0.8059; **Fig. 2g, Supplementary Table 1**). Together, these results suggest trisomic OLLCs are exhibiting moderate differences in migration speed and distance without increasing the relative representation of ventrally-derived OLLCs that migrated into the dorsal compartment of FAs.

### Trisomy drives the enrichment of specific OLLC populations

To characterize the molecular machinery mediating altered OLLC developmental trajectories in trisomy, we profiled 150 *DIV* FAs from three paired isogenic cell lines and two independent differentiations through scRNA-seq and snRNA-seq (**Fig. 3a**). To increase cell capture, due to low OLLC abundance, we combined data from both scRNA-seq and snRNA-seq pipelines, yielding 45,547 cells across all FAs (25,234 euploid and 20,313 trisomic, **Fig. 3b, c**). After integration, we identified 23 distinct clusters (**Supplementary Fig. 3a**) recapitulated in all cell lines, genotypes and sequencing technologies, with quality control metrics assessed across all samples (**Supplementary Fig. 3b–h**), representing major progenitor, neuronal, and glial populations^12,34,45–51^, with broadly similar cell composition between euploid and trisomic FAs (**Fig. 3b–d, Supplementary Fig. 3i, 4, Supplementary Tables 2, 3**).

**Figure 3:**
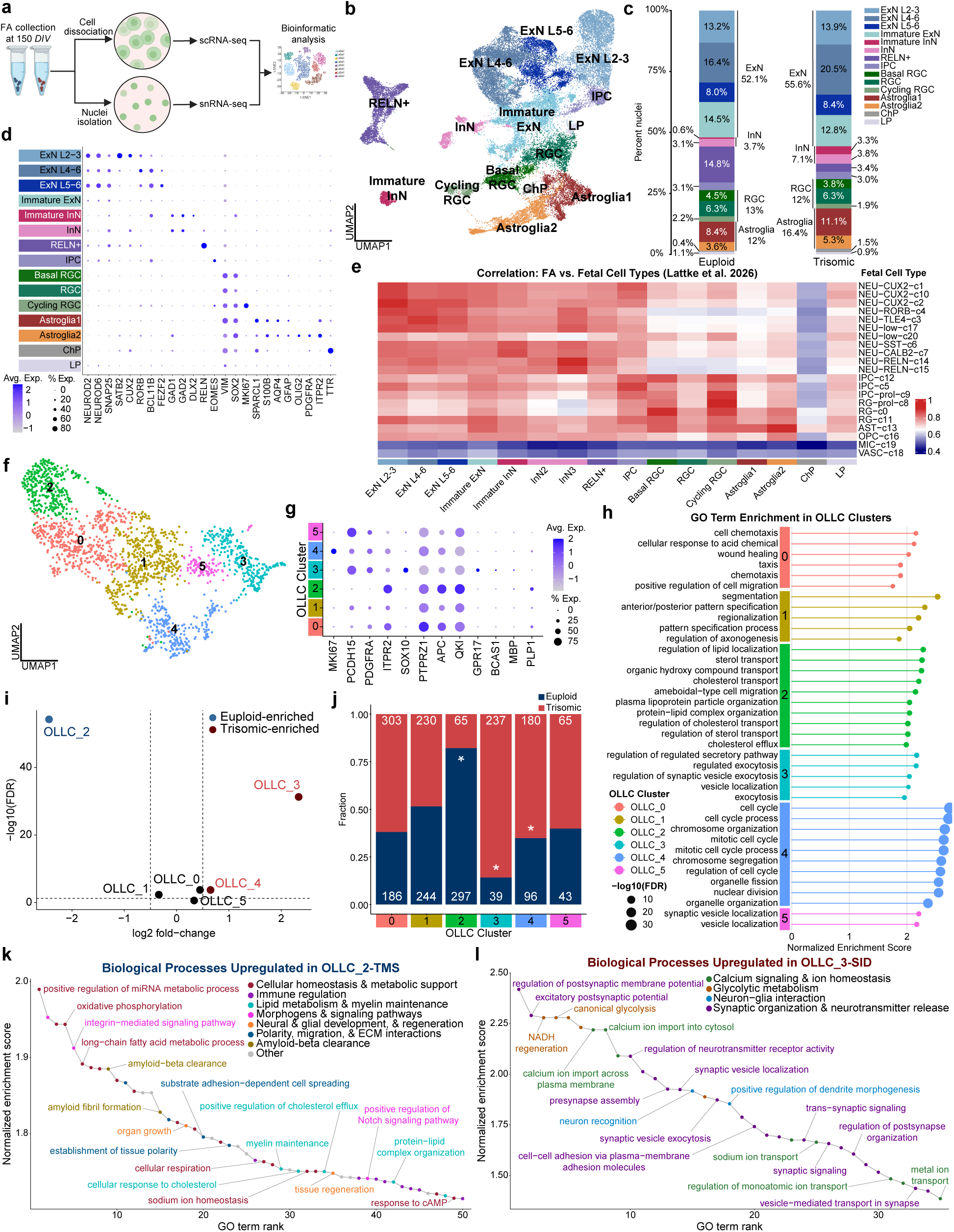
Trisomy induces a transcriptionally distinct OLLC state. **a,** Experimental design and workflow. Forebrain assembloids (FAs) were generated from three pairs of isogenic patient-derived cell lines (euploid and trisomic). Three FAs per genotype from 2 independent differentiation experiments were collected at 150 *DIV* for transcriptional profiling by scRNA-seq and snRNA-seq. Datasets were integrated into a single Seurat object for downstream quality control, normalization, dimensionality reduction, clustering, and visualization. Created in BioRender. Campbell, NB (2026) https://BioRender.com/lklkxmo **b,** Uniform manifold approximation and projection (UMAP) representation of the integrated snRNA-seq and scRNA-seq dataset, displaying cells from all lines, genotypes, and sequencing technologies. Colors indicate major cell types identified by unsupervised clustering of the joint Seurat object and canonical marker expression. **c,** Stacked bar plot showing the relative proportions of major cell populations in euploid and trisomic FAs, including layer-associated excitatory neurons (ExN L2– 3, ExN L4–6, ExN L5–6), immature excitatory neurons (Immature ExN), inhibitory neurons (InN and Immature InN), RELN+ cells, intermediate progenitor cells (IPC), radial glial populations (RGC, basal RGC, cycling RGC), astroglial populations (Astroglia1 and Astroglia2), choroid plexus cells (ChP), and late progenitors (LP). Immature ExNs were identified by excitatory neuronal markers without a defined cortical layer signature, whereas immature InNs expressed *DLX2*. Percentages are indicated for each population. Numerical percentages are shown for each segment. **d,** Dot plot showing native Seurat gene expression for canonical markers (summarized in **Supplementary Table 2**) used for annotation of major cell types, including *NEUROD2, NEUROD6, SNAP25,* (ExNs); *SATB2, CUX2* (ExN L2-3); *RORB, BCL11B, FEZF2* (ExN L4-6, ExN L5-6); *GAD1, GAD2, DLX2, RELN* (InN); *EOMES* (IPCs); *VIM, SOX2, MK167* (radial glial cells); *SPARCL1, S100B, AQP4, GFAP* (astrocytes); *OLIG2, PDGFRA, ITPR2* (OLLCs); and *TTR* (ChP). **e,** Heatmap showing Pearson correlation of expression profiles between FA-derived cell types and human fetal cortical cell types from DS and control samples at postconception week (PCW) 10–20^52^. Columns correspond to fetal cell populations, and rows correspond to FA cell populations. Color scale indicates correlation coefficient (r), highlighting the degree of transcriptomic similarity between FA clusters and fetal reference cell types. **f**, UMAP representation of OLLC reclustering. 1,985 cells from the OLLC clusters identified in **Supplementary** Fig. 4e, f (clusters 0, 6, 7) were subset from the integrated object and further subclustered, yielding six transcriptionally distinct OLLC clusters (0–5) used for genotype-enrichment and gene module analyses. **g,** Dot plot showing native Seurat expression of canonical markers across OLLC subclusters. The dot plot includes progenitor and cycling marker, *MKI67*, as well as canonical markers of OLLC development, including *PDGFRA*, *ITPR2*, *SOX10, PCDH15, PTPRZ1, APC, QKI, BCAS1, GPR17, MBP, PLP1*. Dot size indicates the percentage of cells expressing each gene in a cluster, and color represents average scaled expression. **h,** GO Biological Process enrichment in OLLC subclusters. Gene set enrichment analysis (GSEA; clusterProfiler gseGO) was performed for each OLLC subcluster using a one-versus-rest comparison, with genes ranked by average log_2_ fold change. The analysis was restricted to the 1,155 highly variable genes shared across all cell lines and sequencing-technologies. GO Biological Process terms containing 5–500 genes were tested with 10,000 permutations, and only positively enriched terms with Benjamini–Hochberg-adjusted p < 0.05 are shown (up to 10 per cluster). Terms are grouped by cluster (colored bars, OLLC_0–OLLC_5). The x-axis shows the normalized enrichment score (NES), and point size indicates −log_10_(FDR). **i,** Genotype enrichment of OLLC subclusters. For each OLLC cluster (OLLC_0–OLLC_5), counts of trisomic and euploid cells were tabulated from the Seurat metadata, and enrichment was assessed using Fisher’s exact tests with Benjamini–Hochberg correction. The resulting statistics were summarized as log_2_ fold change of the proportions of trisomic versus euploid cells. A volcano plot shows log_2_ fold change of proportions on the x-axis and negative log_10_FDR-adjusted Fisher p-values on the y-axis, with vertical dashed lines at log_2_ fold change −0.5 and 0.5 and a horizontal dashed line at FDR = 0.05. Using this analysis, OLLC_2 was significantly enriched for euploid cells, whereas OLLC_3 and OLLC_4 were significantly enriched for trisomic cells. **j**, Genotype enrichment across OLLC subclusters. Corresponding bar plot showing the proportion of euploid and trisomic cells assigned to each Harmony-derived OLLC cluster. Asterisks indicate clusters significantly enriched for either genotype, based on two-sided Fisher’s exact tests with Benjamini-Hochberg correction for multiple testing. * FDR-adjusted *p* < 0.05. Gene Ontology (GO) biological process (BP) terms enriched among genes **k**, upregulated in the euploid-enriched OLLC_2-TMS state, highlighting processes related to cellular homeostasis, lipid metabolism, myelin maintenance, and extracellular matrix (ECM) interactions, and **l**, upregulated in the trisomic-enriched OLLC_3-SID state, highlighting processes related to calcium signaling, synaptic organization, neuron–glia interactions and glycolytic metabolic.

Within the two astroglial subclusters, Astroglia1 primarily expressed canonical astrocytic markers (*GFAP*, *AQP4*, *SPARCL1)*, while Astroglia2 expressed canonical OLLC markers (*PDGFRA, OLIG2,* and *ITPR2;* **Fig. 3d, Supplementary Table 3**). Comparison of our dataset with a published fetal human cortical single-cell study, spanning 10–20 postconception weeks (PCWs)^52^, confirmed transcriptional correlation between FA-derived and fetal cell populations, including alignment of our Astroglia2 cluster with fetal OPCs (**Fig. 3e**). However, initial clustering did not fully resolve the OLLC lineage within the astroglial compartment (**Fig. 3b, d**). Therefore, we reclustered both astroglial populations into nine subclusters representing cells across cell lines, sequencing technologies, and genotypes, and identified three subclusters expressing OLLC markers (1,985 OLLCs, 905 euploid and 1,080 trisomic; **Supplementary Fig. 5a–f, Supplementary Table 4**).

Differential expression (DE) analysis between euploid and trisomic cells across the entire OLLC population, identified 1,610 differentially expressed genes (DEGs). Gene ontology (GO) enrichment analysis of DEGs downregulated in trisomy showed the enrichment in processes related to glial cell differentiation, cytoplasmic translation, and oxidative phosphorylation (OXPHOS), while DEGs upregulated in trisomy were enriched in GO terms associated with neuron recognition, synapse organization, and modulation of chemical synaptic transmission (**Supplementary Fig. 5g, h, Supplementary Table 5**), demonstrating broad transcriptional alterations in trisomic OLLCs.

To examine how these transcriptional differences drive oligodendrocyte heterogeneity and developmental states, we subsequently re-clustered the OLLC population, resulting in six transcriptionally-distinct OLLC clusters, each representing all cell lines and sequencing technologies (OLLC_0–OLLC_5; **Fig. 3f–h, Supplementary Fig. 6a–c, Supplementary Tables 6, 7**). OLLC clusters differed in oligodendrocyte-lineage and maturation-associated gene expression (**Fig. 3g**), and included migratory (OLLC_0), early developmental (OLLC_1), structural maturation-associated (OLLC_2), neuronal and synaptic interaction-associated (OLLC_3), proliferative (OLLC_4) and synapse-related populations (OLLC_5, **Fig. 3f–h, Supplementary Fig. 6d, e**). Noticeably, OLLC_2 was strongly enriched in euploid cells, whereas OLLC_3 was primarily composed of trisomic cells (**Fig. 3i, j**) prompting us to focus our subsequent analyses on these predominantly euploid- and trisomic-enriched clusters, OLLC_2 and OLLC_3, respectively.

We identified 3,565 DEGs between OLLC_2 and OLLC_3 (713 upregulated and 2,852 downregulated in OLLC_3), using scMetaIntegrator^53^, which controls for cell line and sequencing technology and GO analysis revealed distinct developmental programs (**Fig. 3k, l, Supplementary Table 8**). OLLC_2 was enriched in processes related to extracellular matrix (ECM) interactions, OXPHOS, tissue morphogenesis, developmental signaling pathways, cellular homeostasis, and importantly, lipid and cholesterol metabolism and myelin maintenance (**Fig. 3k**). Combined with elevated expression of canonical genes such as *ITPR2*^12^, *PLP1*^54^ and *QKI*^55^ (**Fig. 3g**), these findings suggest that OLLC_2 is consistent with a trophic, maturation-supportive state (hereafter referred to as OLLC_2-TMS).

In contrast, OLLC_3, showed increased expression of *SOX10*^35^, *OLIG1, OLIG2*^56–59^*, BCAS1*^40^, and *GPR17*^60–62^, consistent with OLLCs progressing through differentiation but remaining at the pre-myelinating stage^40,60,63^ (**Fig. 3g**). OLLC_3 exhibited enrichment in biological processes involved in calcium signaling and ion transport, synaptic organization and neurotransmitter release, as well as glycolytic and NADH metabolic processes (**Fig. 3l**), indicating that OLLC_3 is characterized by an altered metabolic state with enhanced synaptic interactions and differentiation (hereafter referred to as OLLC_3-SID). Thus, these analyses identify two transcriptionally distinct OLLC states that differ in genotype composition, with OLLC_2-TMS enriched in euploid cells and OLLC_3-SID in trisomic cells. Importantly, transcriptional correlations and principal component analysis (PCA) demonstrated greater similarity within each state across genotypes than between states within the same genotype, supporting their identities as distinct transcriptional states rather than genotype-defined populations (**Supplementary Fig. 6f, g**).

### Trisomy redirects differentiating OLLCs away from maturation-supportive programs towards enhanced neuronal coupling

To uncover coordinated transcriptional programs beyond gene expression changes and GO enrichment in individual clusters, we used Celda to identify 40 co-expression modules in the OLLC dataset, of which 21 modules were enriched in the euploid-associated OLLC_2-TMS state and 13 in the trisomy-associated OLLC_3-SID state (**Fig. 4a, b, Supplementary Fig. 7, 8; Supplementary Tables 9, 10**). GO enrichment analysis of individual modules with clusterProfiler allowed us to assign them to broad functional categories based on shared biological annotations^64^ (**Fig. 4c; Supplementary Table 11**). OLLC_2-TMS-enriched modules were associated with structural maturation and axonal recognition (M2, M29, M31, and M32), trophic support and homeostatic functions (M1, M3, M4, M5, M6, M22, and M30), and mitochondrial OXPHOS and metabolic support (M8–M10 and M18–M20) (**Fig. 4c**).

**Figure 4:**
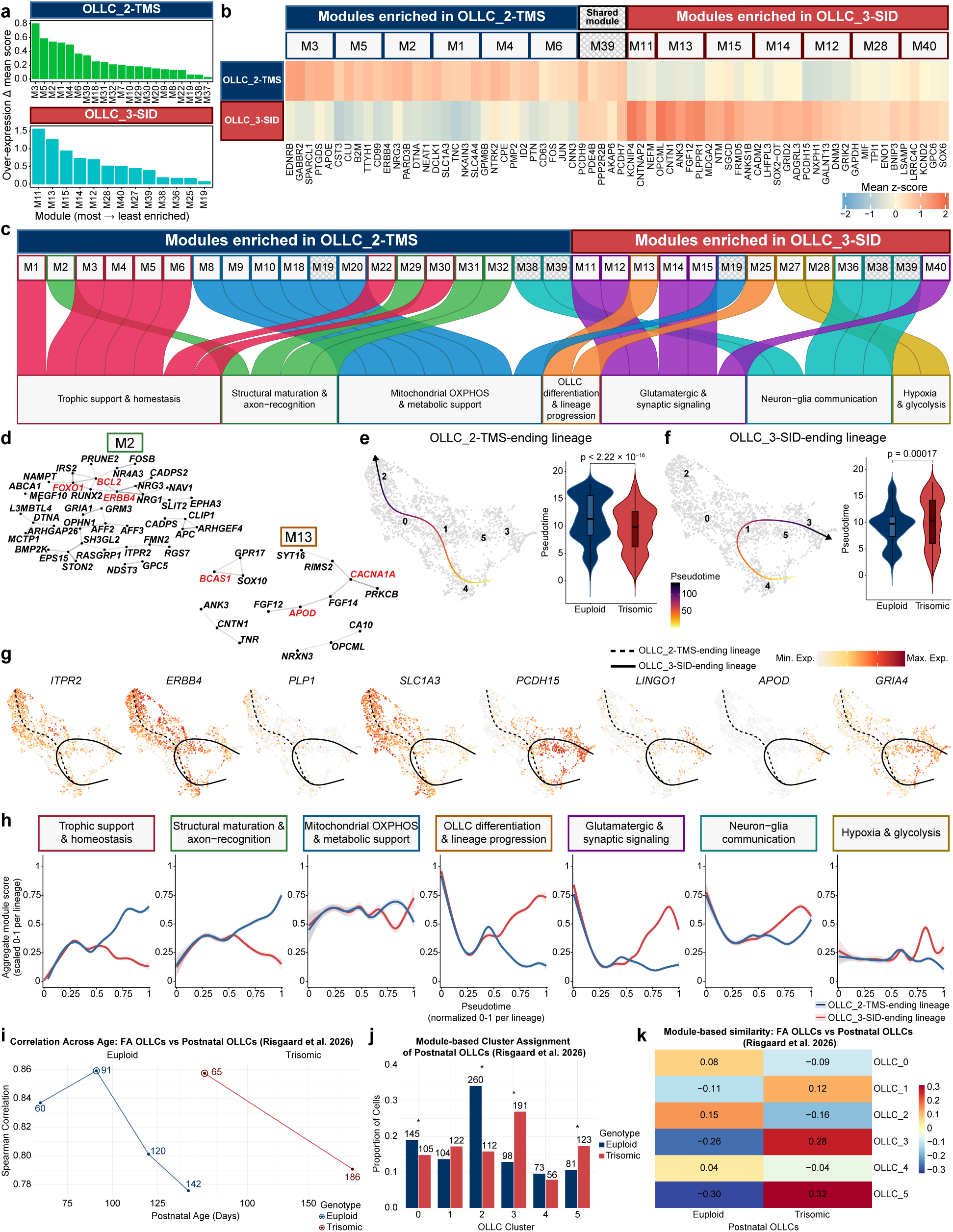
Distinct transcriptional programs define OLLC states and reveal corresponding alterations in the developing human Down syndrome cortex. **a**, Bar plots showing significantly overexpressed Celda gene modules in OLLC_2-TMS and OLLC_3-SID clusters. Celda analysis was used to define a module-feature matrix, and per-cell module scores were computed by z-scoring gene expression across cells and taking the mean z-scored expression of the genes in each module for each cell. For each cluster, Wilcoxon rank-sum tests were performed by comparing module scores in that cluster versus all other OLLCs and p values were adjusted using the Benjamini–Hochberg method. Significantly overexpressed modules were defined as those with FDR < 0.05 and positive mean difference in module score. The plots display all significantly overexpressed modules per cluster, with bar height indicating the extent of change in mean module score. **b,** Heatmap showing the top five expressed genes for overexpressed Celda modules in OLLC_2-TMS and OLLC_3-SID clusters, and shared between the clusters, indicated by crosshatching texture. Rows correspond to OLLC clusters, columns correspond to genes belonging to the overexpressed modules, and values represent scaled expression, highlighting cluster-specific module activity patterns. Modules overexpressed in both clusters are indicated by crosshatching texture. **c,** Alluvial plot showing the distribution of significantly overexpressed Celda modules and their GO-based transcriptional programs in OLLC_2-TMS and OLLC_3-SID clusters. Columns correspond to OLLC clusters, subcolumns indicate individual modules, and bands connect modules to their assigned biological categories at the bottom. Modules overexpressed in both clusters are indicated by crosshatching texture. As such, all modules in the Mitochondrial OXPHOS and metabolic support program were only enriched in OLLC_2-TMS, except M19, which was enriched in both OLLC_2-TMS and OLLC_3-SID. M38 and M39 modules in the Neuron–glia communication were enriched in both OLLC_2-TMS and OLLC_3-SID **d,** STRING association networks of Celda modules M13 and M2. Nodes are module genes, and edges are STRING associations with a combined score ≥ 400; edge transparency reflects the score. Red marks the top three hub genes. **e, f,** Pseudotime trajectories ending in the (**e**) euploid-enriched OLLC_2-TMS cluster and (**f**) trisomic-enriched OLLC_3-SID cluster overlaid on the OLLC UMAP (left), and violin plot showing the distribution of pseudotime values along each lineage for euploid and trisomic cells (right); OLLC_2-TMS-ending lineage: *p* < 2.22 × 10^−16; OLLC_3-SID-ending lineage: *p* = 0.00017 (Wilcoxon rank-sum test). **g,** Trajectory-overlaid feature UMAPs showing expression of selected genes (*ERBB4, ITPR2, PLP1, SLC1A3, PCDH15, LINGO1, APOD, GRIA4*) along the two pseudotime lineages of interest. Cells are colored by normalized expression (from minimum to maximum), with the OLLC_2-TMS-ending lineage shown as a dashed curve and the OLLC_3-SID-ending lineage shown as a solid curve. **h**, Fate-conflict plots showing aggregate module scores for GO-based gene programs across pseudotime in the OLLC_2-TMS-ending (blue) and OLLC_3-SID-ending (red) lineages. Each panel corresponds to one gene program, with the y-axis indicating lineage-scaled aggregate module scores and the x-axis indicating normalized pseudotime (0–1 per lineage). Shaded area represents 95% confidence intervals. **i,** Spearman correlation between FA OLLCs and postnatal OLLCs at each available postnatal age, calculated across all shared genes, to identify the postnatal timepoint that most closely aligns with the FA OLLCs. Euploid (days 60, 91, 120 and 142) and trisomic (days 65 and 186) samples are shown separately, and point labels give the age in days. Circled points indicate the highest correlation with FA OLLCs per genotype. **j,** Proportion of postnatal euploid (blue; n = 761) and trisomic (red; n = 709) cells assigned to each OLLC subcluster by module-based scoring. Numbers above the bars show cell counts. *BH FDR < 0.05, Fisher’s exact test. **k,** Heatmap of mean similarity scores for each OLLC subcluster (rows) in postnatal euploid (day 60) and trisomic (day 65) cells (columns). Positive values (red) indicate higher-than-average expression of a signature, negative values (blue) lower-than-average, and values near zero no difference.

Structural maturation and axon-recognition programs contained markers of oligodendrocyte lineage-commitment and maturation, axon-recognition, and myelin maintenance, including *ITPR2*^47,65^*, APC*^66^, *PTPRZ1*^67^, *ERBB4*^68–70^, *NRG3*^71^, *NRCAM*^72,73^, *QKI*^55^, and *RTN4*^74^, indicating a structurally mature OLLC state, poised for axonal ensheathment and myelination. Consistent with this interpretation, STRING protein-association analysis of a representative module (M2) identified functional networks associated with axon–glia interactions, including ERBB4–NRG1/NRG3 signaling (**Fig. 4d**).

Homeostasis and trophic support modules included genes involved in glutamate uptake and recycling, including *SLC1A3*^75^ and *GLUL*^76^, lipid homeostasis, including *PTGDS*^77^, *APOE*^78^, *CLU*^79^, and trophic and growth-related signaling, including *TGFB2*^80^, *PTN*^81^, *TNC*^82^, *MDK*^83^, and *NTRK2*^84^. Mitochondrial OXPHOS-related modules contained genes encoding for mitochondrial respiration chain and ATP production machinery, such as *ND1–ND6*^85^, *COX1–3*^86–88^*, ATP8, COX4I1*^85^ and *PINK1*^89^, reflecting increased bioenergetic needs required for membrane and process extension and myelination^86,90^, suggesting that OLLC_2-TMS state participates in coordinated trophic, structural and metabolic programs supporting productive maturation.

In contrast, OLLC_3-SID-enriched modules were associated with oligodendrocyte differentiation and lineage progression (M13 and M25), glutamatergic and synaptic signaling (M11, M12, M14, M15, and M40), neuron–glia communication (M36, M38, and M39) and hypoxia-related processes (M27, M28; **Fig. 4c**). Differentiation and lineage progression-associated modules contained drivers of oligodendrocyte differentiation, *OLIG1*, *OLIG2*^56–59^, and *BCAS1*^35,40^, and *CNTN1* and *ANK3*, which regulate oligodendrocyte–neuron interactions^91,92^. STRING analysis of M13 identified predicted functional associations among *BCAS1*, *SOX10*, and *GPR17*, and a distinct *CNTN1–ANK3–TNR* group associated with axon–glia interactions (**Fig. 4d**). Notably, GPR17, a "cell-intrinsic timer" of oligodendrocyte differentiation, which is downregulated in myelinating oligodendrocytes^60,63^, was co-expressed with BCAS1, without corresponding enrichment in myelin structural programs, indicating that trisomic OLLCs reach the pre-myelinating stage without progressing to myelination, mirroring the expansion of BCAS1+/MBP− cells observed in CC1+ OLLCs in trisomic FAs (**Fig. 1i, l**).

The glutamatergic and synaptic signaling modules contained genes that mediate adhesion and synaptic scaffolding such as *CADM2*^93^, *MDGA2*^94,95^, *LRRTM4*^96^, *LHFPL3*^97^, and *DLGAP1*^96^, and genes mediating glutamatergic signaling and excitability, including *GRIA4*^98–100^, *GRIA2*^99^, *CNTNAP2*^101,102^, and *KCNIP4*^101^. Notably, M14 also contained *LINGO1*, a negative regulator of myelination^103^. Taken together, these findings reveal enhanced synaptic engagement in OLLC_3-SID, while enrichment in *LINGO1* and *GPR17* indicates two independent brakes on the transition from pre-myelinating to myelinating OLLCs.

Neuron–glia communication modules M38 and M39 were shared between the two states, while M36 was additionally enriched in OLLC_3-SID (**Fig. 4a, c**). These modules contained genes encoding for cell adhesion and signaling molecules such as *NCAM2*^104^, *PCDH7*, *PCDH9*^105^, and *PTPRT*^106,107^, a regulator of neurite outgrowth and axon guidance, *SEMA6D*^108^, genes regulating excitability, including *AKAP6/9*^109^ and *SHISA6*^110^, and an inhibitory postsynaptic scaffold protein, *GPHN*^111^, suggesting the response to synaptic signaling is coordinated and preserved across OLLC states. The hypoxia and glycolysis-related program contained canonical glycolytic enzymes *GAPDH*, *TPI1,* and *ENO1*^112^, further distinguishing OLLC_3-SID. Overall, these results indicate a state-specific signature of pre-myelinating OLLCs with increased engagement in neuronal and synaptic signaling, and glycolytic processes that lack the expected coordination with myelination-supportive processes, consistent with the increased proportion of CC1+/BCAS1+/MBP-OLLCs observed in trisomic FAs (**Fig. 1i, l**).

To determine how these transcriptional signatures diverge during oligodendrocyte lineage progression, we reconstructed developmental trajectories using pseudotime^113^. Using the shared population of cycling progenitors (OLLC_4) as the root state^114,115^, we identified two major trajectories terminating in the OLLC_2-TMS and OLLC_3-SID clusters (**Fig. 4e, f**). Trisomic OLLCs were enriched at earlier pseudotime along the OLLC_2-TMS-ending trajectory and later pseudotime along the OLLC_3-SID-ending trajectory compared to euploid OLLCs, indicating OLLCs from the two genotypes preferentially occupy different positions along these two lineages. Genes associated with oligodendrocyte structural support, oligodendrocyte-axonal interaction and myelination potential, including *ITPR2*^47,54^*, ERBB4*^68,70^, *PLP1*^116^ and *SLC1A3*^75^, were preferentially expressed along the OLLC_2-TMS-ending trajectory. In contrast, *PCDH15*^117^, *APOD*^28,118^, *LINGO1*^119^, and the AMPA receptor subunit GRIA4^99^ were associated with OLLC_3-SID-ending trajectory (**Fig. 4g**). In line with this, aggregate module expression of trophic support, homeostasis, structural maturation, and axon-recognition programs increased along the OLLC_2-TMS-ending lineage, while expression levels were maintained or declined along the trisomic OLLC_3-SID-ending lineage. Conversely, oligodendrocyte differentiation, glutamatergic/synaptic signaling, and hypoxia/glycolysis programs increased preferentially along the OLLC_3-SID-ending lineage, and mitochondrial OXPHOS and neuron–glia communication programs demonstrated similar patterns between the trajectories (**Fig. 4h**).

Since myelination in the prefrontal cortex begins late in gestation (around 32GW), and primarily occurs postnatally^120,121^, we examined whether these transcriptional states are also represented in developing postnatal human DS brains. We projected FA-derived OLLC signatures onto published postnatal DS prefrontal cortex transcriptomic data^114^. Pseudobulk correlations supported the selection of euploid day 60 and trisomic day 65 OLLCs for reference mapping (**Fig. 4i** and module-based mapping of human OLLCs onto FA OLLC clusters revealed preferential assignment of euploid cells to OLLC_2-TMS and trisomic cells to OLLC_3-SID (**Fig. 4j, k**). Together, these findings indicate that trisomic OLLCs follow a differentiation trajectory towards a pre-myelinating state with increased synaptic engagement, lacking complementary enrichment in myelination-related signatures, which is consistent with OLLC states found in early postnatal human DS cortex.

### OLLC_2-TMS and OLLC_3-SID exhibit distinct patterns of bidirectional communication with neurons

To determine whether these transcriptional states differ in their predicted interactions with neurons, we used CellChat to infer bidirectional ligand–receptor (L–R) signaling between the OLLC states and mature and immature excitatory (ExNs) and inhibitory neurons (InNs; **Fig. 5a, b**). Both states communicated most extensively with mature neurons, while OLLC_2-TMS exhibited more numerous and stronger interactions with both ExNs and InNs, particularly with mature ExNs. The differences were more pronounced in reciprocal neuron-to-OLLC communication, with OLLC_2-TMS receiving stronger and more extensive predicted inputs from mature ExNs and InNs (**Supplementary Fig. 9a, b**). Despite enrichment in neuronal-interaction programs (**Fig. 4a–c, g, h**), OLLC_3-SID did not show global enhancement in neuron–glia communication. This apparent discrepancy suggests that OLLC_3-SID may engage in selective, pathway-specific interactions, prompting investigation into underlying signaling pathways.

**Figure 5:**
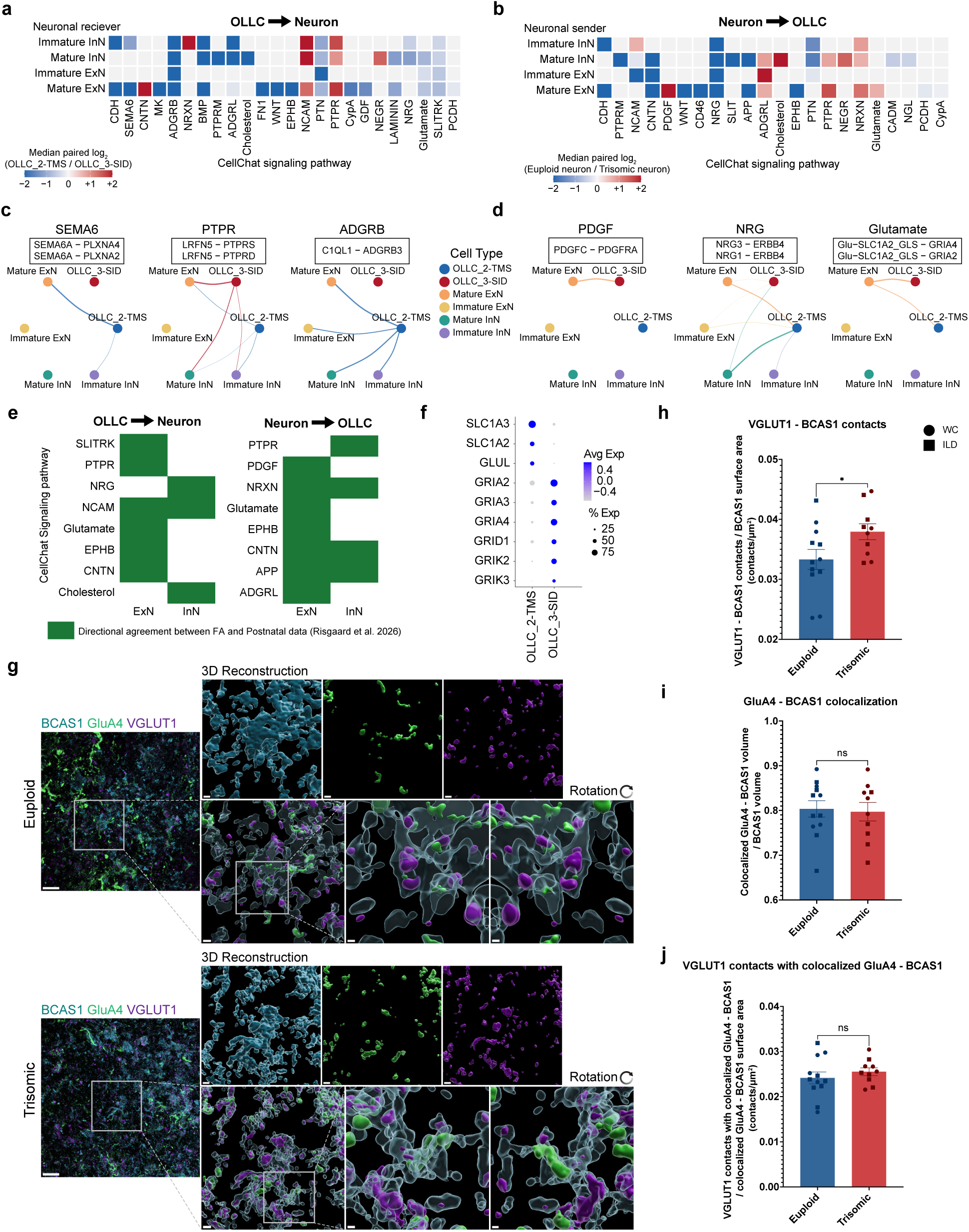
Trisomy 21 is associated with reorganization of neuron–OLLC communication. **a, b,** Consensus heatmaps of CellChat signaling pathways for (**a**) OLLC-to-neuron communication, with neuronal populations as receivers, and (**b**) neuron-to-OLLC communication, with neuronal populations as senders. Communication involving OLLC_2-TMS was inferred in euploid samples and communication involving OLLC_3-SID in trisomic samples. For each pathway and neuronal population, paired log_2_ fold changes in pathway probability were calculated within six matched cell line × sequencing-technology pairs and summarized as the median. Tiles are colored only for combinations with a consistent direction of change in at least three matched pairs; grey tiles indicate no consensus. Blue, higher inferred signaling involving OLLC_2-TMS; red, higher inferred signaling involving OLLC_3-SID; color scale capped at ± 2. **c, d,** Consensus circle plots for selected pathways: (**c**) OLLC-to-neuron signaling (SEMA6, PTPR, ADGRB) and (**d**) neuron-to-OLLC signaling (PDGF, NRG, Glutamate). Nodes represent OLLC_2-TMS, OLLC_3-SID and neuronal populations (mature and immature ExNs and InNs). Edge width represents the median inferred pathway probability across matched pairs, and edges are colored by sender population. **e**, Heatmaps showing concordance of predicted neuron–OLLC signaling pathways between FAs and early postnatal human prefrontal cortex^114^. Green indicates pathways exhibiting consistent directional preferences across datasets, stratified by signaling direction (OLLC to neuron and neuron to OLLC) and neuronal population (ExN and InN). **f,** Dot plot showing expression of glutamate transporters (*SLC1A3*, *SLC1A2*), glutamine synthetase (*GLUL*), and ionotropic glutamate receptor subunits (*GRIA2*, *GRIA3*, *GRIA4*, *GRID1*, *GRIK2*, and *GRIK3*) in OLLC_2-TMS and OLLC_3-SID. Color indicates average scaled expression and dot size the percentage of cells expressing each gene. **g,** High resolution confocal representative images and corresponding reconstructions of BCAS1, GluA4, and VGLUT1 expression in trisomic and euploid FAs at 150 *DIV*. Rotated reconstructions show GluA4-BCAS1 colocalization and contacts between VGLUT1, GluA4, and BCAS1. Scale bars, 20 µm (confocal images), 3 µm (reconstructions), and 1 µm (reconstruction insets and rotations). **h–j,** Quantification of (**h**) VGLUT1-BCAS1 contacts (defined as VGLUT1 expression within 1 µm of BCAS1 expression) normalized to BCAS1 surface area (contacts/µm^2^; LME, β = + 0.0046*, p* = 0.03796), (**i**) Colocalized GluA4-BCAS1 volume (defined as overlapping expression of both proteins with a volume > 0.01 µm^3^) normalized to BCAS1 volume (LME, β = -0.0085*, p* = 0.6800), and (**j**) VGLUT1 contacts with colocalized GluA4-BCAS1 (defined as VGLUT1 expression within 1 µm of colocalized GRIA4-BCAS1 expression) normalized to GluA4-BCAS1 surface area (contacts/µm^2^; LME, β = +0.0014*, p* = 0.3902) per FA. Euploid, *n* = 12. Trisomic, *n* = 10 FAs. Graphs display results from two independent differentiation experiments. All comparisons were analyzed using linear mixed-effects models, with genotype effect reported in the figure legend and the effects of cell line and differentiation summarized in **Supplementary Table 1**. Each point represents one FA and each symbol shape represents an isogenic cell line pair. circle = WC; square = ILD. Results shown as mean ± SEM. * *p* < 0.05, ns = not significant.

To identify state-specific signaling pathways, we retained those showing concordant differences in inferred CellChat communication probabilities across at least three matched OLLC_2-TMS and OLLC_3-SID comparisons from three isogenic pairs (**Fig. 5a, b; Supplementary Fig. 9c, d**). Several homophilic adhesion pathways with limited directional interpretation were preferentially associated with OLLC_2-TMS–neuron interactions (CDH^122^ and PTPRM^123^) or OLLC_3-SID–neuronal interactions (NEGR^124^; **Fig. 5a, b; Supplementary Fig. 10, 11**).

Among directional pathways, OLLC_2-TMS preferentially communicated with neurons via trophic and pro-survival signaling (PTN^81^ and MK^125,126^ pathways) and adhesion and axon-recognition signaling pathways (SEMA6^127^ and ADGRB^128^; **Fig. 5a, c**). In contrast, OLLC_3-SID preferentially engaged in synaptic adhesion and organization pathways, including the NRXN^101^, and PTPR^129,130^. Both OLLC states engaged in PTPR signaling, mediated by LRFN5–PTPRD, with stronger communication from OLLC_3-SID to mature ExNs and InNs (**Fig. 5a, c; Supplementary Fig. 10**). PTPRD participates in synaptic assembly and is transiently upregulated in maturing oligodendrocytes, where it can regulate initiation of myelination^129,130^.

Reciprocal neuron-to-OLLC signaling favored trophic and axon–glia pathways in OLLC_2-TMS including NRG signaling, which has been implicated in peripheral and central nervous system myelination^131,132^, primarily mediated by NRG3–ERBB4 across neuronal populations (**Fig. 5b, d**). In contrast, OLLC_3-SID preferentially engaged in PDGF signaling, mediated by PDGFC– PDGFRA, regulating OLLC development, migration, and survival^133^, and neuronal and synaptic-associated glutamatergic signaling, particularly from mature ExNs (**Fig. 5b, d, Supplementary Fig. 11**).

To determine if these signaling patterns were conserved in postmortem human samples, we performed CellChat analysis on the postnatal DS prefrontal cortex dataset^114^, comparing human euploid OLLCs assigned to OLLC_2-TMS state via module-based mapping to human trisomic OLLCs corresponding to OLLC_3-SID state (**Fig. 4j**). Remarkably, several pathways identified in our FAs showed concordant predicted signaling with the postnatal dataset, supporting these state-specific signaling patterns in Trisomy 21 (**Fig. 5e**). The preferential glutamatergic signaling involving OLLC_3-SID indicated altered interactions between trisomic OLLCs and neuronal synaptic compartment. Consistent with this, OLLC_3-SID showed higher expression of glutamate receptor genes (*GRIA2*, *GRIA3*, *GRIA4*, *GRID1*, *GRIK2*, *GRIK3*) while OLLC_2-TMS showed higher expression of glutamate transporters (*SLC1A3*, *SLC1A2*) and *GLUL* (**Fig. 5f**), suggesting state-related differences in glutamate processing.

To evaluate interactions between ExNs and OLLCs at the protein level, we quantified contacts and colocalization between immunolabeled CC1+ and BCAS1+ OLLCs, presynaptic terminal marker VGLUT1, and postsynaptic AMPA receptor subunit, GluA4, encoded by *GRIA4* (**Fig. 5g**). Trisomic FAs exhibited an increased number of VGLUT1 contacts with BCAS1+ OLLCs (*p=*0.03796; **Fig. 5g, h, Supplementary Table 1**), suggesting enhanced ExN–glia coupling in trisomic pre-myelinating OLLCs. However, we did not detect differences in GluA4-BCAS1 colocalized volume (*p=*0.6800; **Fig. 5g, i**), or in the number of VGLUT1 contacts with colocalized GluA4-BCAS1 (*p=*0.3902; **Fig. 5g, j**) in trisomic FAs. Furthermore, neither the postsynaptic scaffolding protein, HOMER1b/c colocalization with BCAS1 (*p=*0.0.1448), nor PSD95-BCAS1 colocalization (*p*=0.3413) differed significantly between the genotypes (**Supplementary Fig. 12a–d**). These results suggest that the enhanced glutamatergic signaling predicted by CellChat aligns with increased VGLUT1 contacts with BCAS1+ OLLCs, but does not result in increased GluA4 or postsynaptic scaffolding protein expression in trisomic BCAS1+ OLLCs.

In contrast to the increased contacts between VGLUT1 and BCAS1+ OLLCs, contacts between VGLUT1 and CC1+ OLLCs did not differ significantly (*p=*0.5653; **Supplementary Fig. 12e, f**), indicating that altered excitatory neuron–glia coupling might be developmental stage specific. Lastly, we found no difference in the number of VGAT-BCAS1 contacts (*p=*0.8535; **Supplementary Fig. 12g, h**), revealing that the increase in excitatory presynaptic contacts with BCAS1+ pre-myelinating OLLCs in trisomic FAs was not accompanied by an equivalent increase in inhibitory presynaptic contacts.

### Trisomic FAs contain altered extracellular compact myelin and increased frequency of defective myelin

To evaluate how the shift in the developmental trajectory of trisomic OLLCs affects myelination, we examined myelin ultrastructure with transmission electron microscopy (TEM) in our FAs at 150 *DIV*. We observed myelin in both the intracellular and extracellular spaces, and classified myelinated axons based on the level of compaction (**Fig. 6a, b**). Interestingly, the majority of myelinated axons were intracellular, and axons surrounding these structures were mostly unmyelinated in both genotypes. Quantitative analyses of identified myelinated axons using TEM did not show significant differences in the number of myelinated axons, the proportion of intra-/extracellular myelin or proportions of loose, mid-compact, and compact myelin between our euploid and trisomic FAs (**Fig. 6a–e, Supplementary Fig. 13a, b**). We then assessed the myelin thickness in the mid-compact and compact groups. In trisomic FAs, we found significantly thinner myelin sheaths in the extracellular compact group (*p*=0.0437), with 100% of our trisomic samples falling into the thin group (thickness < 0.5006 µm; **Fig. 6a, f, Supplementary Fig. 13c– e**). We also observed significantly increased numbers of defective myelin sheaths in our trisomic FAs, classified by the presence of either detachment of myelin sheaths from the axon, dark axoplasm, irregular lamellar splitting or redundant myelin formation (*p*=0.0065; **Fig. 6g, h Supplementary Figure 13 f, g**). Complementary immunostaining of FAs with BCAS1, MBP, and NF-H, which marks neuronal processes, showed that MBP was predominantly colocalized with BCAS1+ processes, and NF-H+ axons were mostly unmyelinated (**Fig. 6i**). Consistent with our molecular data, these data suggest dysregulated axonal myelination dynamics in trisomic FAs.

**Figure 6:**
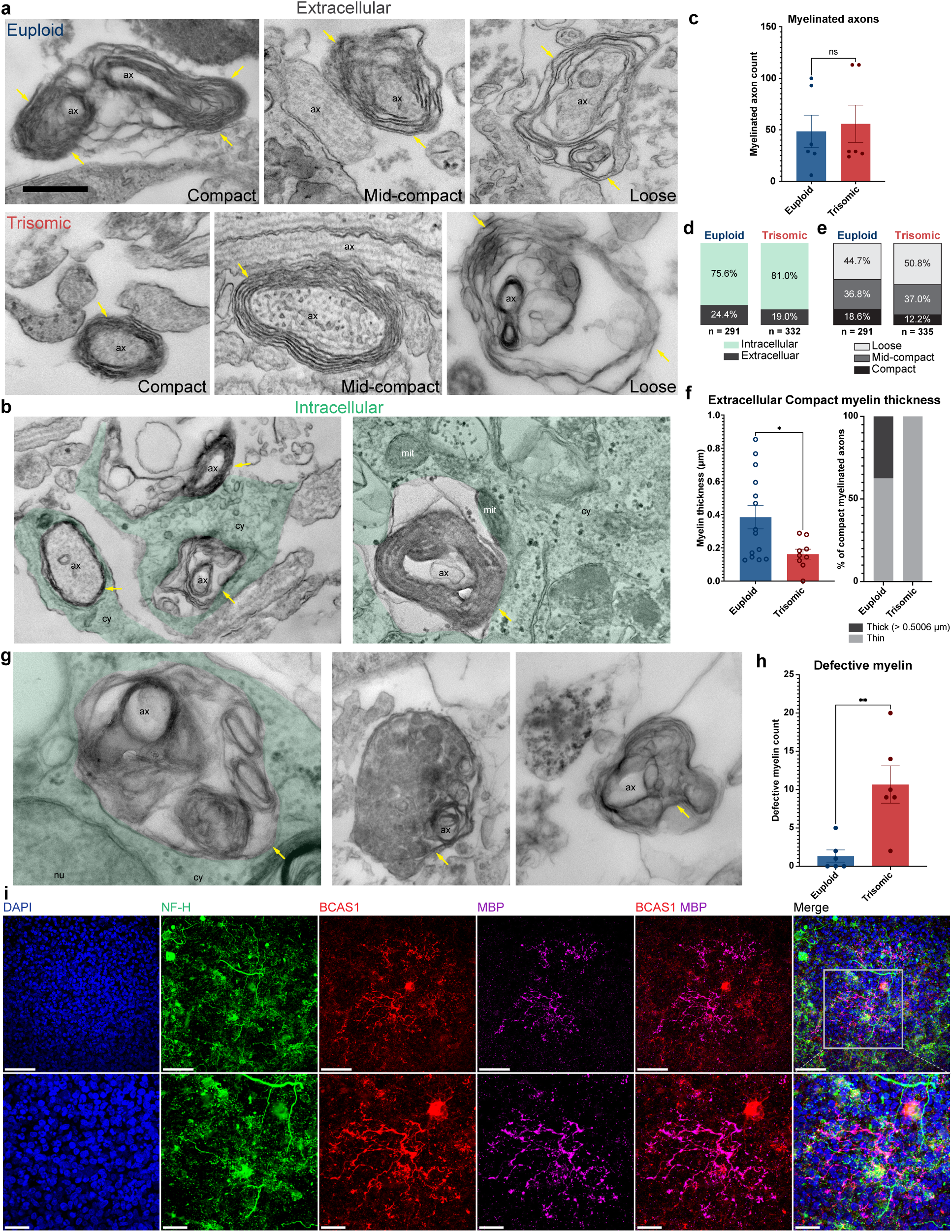
Trisomic FAs exhibit abnormal myelin phenotypes. **a**, **b,** Electron micrographs showing (**a**) loose, mid-compact, and compact extracellular myelin in euploid and trisomic FAs at 150 *DIV* and (**b**) intracellular myelin, surrounded by cytoplasm (green). ax, axon; cy, cytoplasm (green); mit, mitochondria; nu, nucleus. Yellow arrows indicate myelin. **c,** Quantification of myelinated axon count (Mann Whitney test, *p* = 0.8355) per FA. Each point represents one FA. **d, e,** Stacked bar graphs showing (**d**) intracellular (green) and extracellular (dark grey) myelin and (**e**) loose (light grey), mid-compact (dark grey) and compact (black) myelin proportions in FAs. **f,** Quantification of extracellular compact myelin thickness (µm; Mann Whitney test, *p* = 0.0437). Each open circle represents one myelinated axon. Stacked bar plot showing extracellular thick (dark grey) and thin (light grey) compact myelin proportions. Thick myelin is defined as myelin thicker than 0.5006 µm. **g,** Electron micrographs showing defective myelin at 150 *DIV*. Yellow arrows indicate defects such as splitting between myelin layers. **h,** Quantification of defective myelin (Mann Whitney test, *p* = 0.0065) per FA. Each point represents one FA. Euploid, *n* = 6. Trisomic, *n* = 6 FAs. **i**, Representative stitched and inset confocal images showing NF-H (green), BCAS1 (red), and MBP (magenta) expression from one FA at 150 *DIV*. Scale bars, 50 µm and 20 µm (insets). Graphs display results from two independent differentiation experiments. Results shown as mean ± SEM (**a, h**), proportions (**d–f**), and sums (**f**). * *p* < 0.05, ** *p* < 0.01, ns = not significant.

## Discussion

In the current study, we utilized an *in vitro* human FA model to examine the effect of Trisomy 21 on oligodendrocyte lineage development within a multicellular system. Our findings reflect consistent phenotypes identified across three isogenic paired cell lines derived from individuals with DS. Rather than delaying OLLC maturation, trisomy preferentially expands the population of pre-myelinating OLLCs. These findings suggest that the myelination deficits previously reported in DS mouse models and human tissue^2,5,6,114,134,135^ may arise, at least in part, from a disrupted coordination between oligodendrocyte differentiation and the maturation programs supporting productive myelination.

This interpretation is supported by several complementary findings. First, trisomic FAs contained more CC1+/BCAS1+ OLLCs, including more CC1+/BCAS1+/MBP− cells. Second, trisomic OLLCs preferentially adopted the OLLC_3-SID state, marked by neuronal and synaptic programs and lacking programs supporting myelination present in the euploid-enriched OLLC_2-TMS state, revealing that trisomy shifts OLLCs away from OLLC_2-TMS and toward OLLC_3-SID. Third, trisomic FAs showed an increase in defective myelin ultrastructure, and thinner extracellular compact myelin sheaths.

Alterations in OLLC development were also evident in their migratory behavior. Live-cell imaging showed that trisomic *SOX10*+ OLLCs migrated at higher speeds and over greater distances than their euploid counterparts. However, these differences were not reflected in the overall number of migrating cells reaching the dCO portion of FA, suggesting that trisomy does not impair the overall capacity of OLLCs to populate the developing forebrain, but rather alters the dynamics of their migration. These findings may be related to elevated *PCDH15* expression in OLLC_3-SID (**Fig. 4g**), which promotes motility in OPCs^117^ or to the overexpression of *OLIG2,* which has been shown to enhance the migratory capacity of OPCs in a transgenic mouse model^136^, raising the possibility that *OLIG2* triplication in DS could be involved in the increased motility of trisomic OPCs. Alternatively, these differences may reflect altered interactions with the surrounding neuronal environment, potentially involving the dysregulated glutamate signaling identified in trisomic OLLCs. Future studies will be required to distinguish between OLLC-intrinsic alterations and altered environmental cues contributing to their migratory behavior.

Transcriptomic analysis identified two OLLC states with distinctive differentiation, maturation, and neuron–glia interaction programs. The euploid-enriched OLLC_2-TMS state demonstrated coordinated maturation with trophic, metabolic, structural, and axon-recognition programs required to support myelination, and included the upregulation of established regulators of oligodendrocyte development and myelination, such as *QKI*^55,137^*, ITPR2*^47,65^*, APC*^66^*, PTPRZ*^67^, *SEMA6A*^127^ and components of *NRG/ERBB* signaling^68,70^. These transcriptional features were accompanied by extensive predicted reciprocal neuron–OLLC communication through trophic, ECM, adhesion, and axon-interaction pathways, mediated by PTN, SEMA6, NRG, LAMININ, CDH, PTPRM, EPHB, CNTN, and ADGRB. Notably, predicted C1QL1–ADGRB3 and SEMA6A– PLXNA2 interactions are particularly relevant to myelination. C1QL1, expressed predominantly by OPCs, plays a critical role in promoting myelination and remyelination^138,139^, while SEMA6A expression peaks during active myelination and its loss inhibits oligodendrocyte maturation and myelin production^127^. Thus, OLLC_2-TMS appears to represent a state in which oligodendrocyte maturation is coordinated with the acquisition of trophic-supportive and axonal ensheathment functions.

In contrast, OLLC_3-SID showed upregulation of markers of oligodendrocyte differentiation, including *SOX10*, *BCAS1*^40^, and *APOD*^28^. The co-expression of these markers alongside sustained upregulation of *GPR17*^62,140^ and myelination inhibitor *LINGO1*^119^ supports a state that has progressed through differentiation, but remains unable to transition. OLLC_3-SID was dominated by glycolytic programs, whereas OLLC_2-TMS displayed upregulated OXPHOS and lipid and cholesterol metabolism programs that are necessary to meet the biosynthetic demands of process extension and myelination^86,90^. These differences suggest that OLLC_3-SID represents a distinct differentiation state, in which differentiation-associated and synaptic programs advance while programs required for myelination do not. Previous studies in Ts65Dn mice have demonstrated incomplete oligodendrocyte maturation despite the acquisition of differentiation markers, ultimately resulting in hypomyelination^6^. Our findings extend these observations to human trisomic OLLCs and identify distinct transcriptional programs that may underlie this divergence between differentiation and productive myelination.

Interestingly, the overrepresented neuronal-interaction profile of OLLC_3-SID did not correspond to a global increase in communication with neurons. Instead, it showed fewer and weaker predicted interactions with mature ExNs and InNs and was selectively enriched in neuronal recognition and synaptic-associated pathways, including NRXN, NCAM, PTPR, and CNTN. This signaling reorganization was also observed in reciprocal neuronal input to OLLCs, where NRXN and, notably, glutamatergic signaling dominated the pathways leading from mature ExNs to OLLC_3-SID. This suggests that OLLC_3-SID–neuronal communication is reorganized, rather than amplified and shifted toward more selective neuronal and synaptic-associated signaling.

Importantly, the OLLC state divergence identified in our FAs was also evident in the developing human DS brain. A recent single-nucleus multiomic analysis of postnatal DS prefrontal cortex reported altered oligodendrocyte lineage progression across samples spanning 2 months to 3 years of age^114^. This period encompasses substantial changes in the oligodendrocyte lineage, with OPC numbers peaking at birth, and progressive myelination throughout the first two years of life^120,121^. Although brain organoids show transcriptional alignment with midgestational human brain^141,142^ (**Fig. 2e**), this transcriptional global correspondence does not necessarily represent the developmental stage of OLLCs in our FAs. In the human brain, myelination progresses in a caudal-to-rostral direction, with the forebrain among the latest regions to myelinate^121^. In fact, at 20 GW, myelinated axons are present in the diencephalon but absent from the telencephalon, with cerebral white matter myelination beginning at approximately 32 GW^120,143^. Thus, the presence of pre-myelinating and myelinating OLLCs in our FAs justifies the comparison with a later developmental window.

Consistent with this, FA-derived OLLCs showed stronger alignment with early (day 60/65) human postnatal OLLC populations^114^. Remarkably, human OLLCs showed genotype-biased representation across these states, with trisomic cells depleted from the maturation-supportive state and enriched in a restricted pre-myelinating state associated with neuronal and synaptic programs. This correspondence indicates that OLLC_2-TMS and OLLC_3-SID capture biologically relevant OLLC states present in the developing human DS cortex rather than FA-specific transcriptional phenotypes. Notably, oligodendrocyte development in the human brain includes a prolonged period, beginning around 30 GW, associated with the presence of immature oligodendrocytes that have differentiated but have not yet started active myelination^144^. Therefore, the enrichment of trisomic cells in OLLC_3-SID state in both systems raises the possibility that trisomy prolongs this normally transient pre-myelinating state, thus inflating the population of early pre-myelinating OLLCs in trisomy.

This biological alignment included predicted neuron-to-OLLC communication, with several state specific pathways in FAs showing corresponding state-associated differences in the day 60–65 human cortex, including (NRXN, PDGF, and glutamate pathways). Together, the recapitulation of both OLLC states and signaling pathways support the identified split in the developmental trajectory in our FA model.

Consistent with the enrichment of glutamatergic signaling transcriptomic signatures in FAs and human datasets, trisomic OLLCs also showed increased association with excitatory neuronal synaptic markers, whereas VGAT+ contacts did not differ between genotypes. These data suggest that altered engagement of trisomic OLLCs preferentially involves ExNs rather than broadly increased neuronal interactions.

Previous studies suggest that the transition from OPCs toward myelinating oligodendrocytes involves reduced synaptic input and glutamatergic signaling, whereas persistent neuronal engagement may interfere with oligodendrocyte maturation and axonal ensheathment^10,15^. Consistent with this, OLLC_2-TMS demonstrated increased expression of glutamate transporter *SLC1A3* and glutamine synthetase *GLUL*, indicating an increased capacity for glutamate handling, and correspondingly low expression of ionotropic glutamate receptors. In contrast, OLLC_3-SID showed a reciprocal profile with elevated AMPA receptor subunit transcript expression. Despite increased VGLUT1 contacts with BCAS1+ OLLCs in trisomic FAs, no corresponding increase was detected in colocalization of BCAS1+ cells with GluA4 or postsynaptic scaffolding proteins, HOMER1b/c and PSD95, suggesting that enhanced glutamatergic synaptic engagement in trisomic OLLCs is not reflected in postsynaptic structural organization, required for myelin sheath formation^93^. Together, these findings highlight a divergence between glutamate handling and glutamatergic receptor-associated programs between OLLC states, alongside increased excitatory neuronal engagement in trisomic OLLCs without a corresponding enhancement in postsynaptic assembly.

The present study does not establish whether increased glutamatergic signaling causatively contributes to the establishment of OLLC_3-SID state or the abnormal myelin formation. Although altered neuronal activity has previously been reported in trisomic iPSC-derived neurons^19^, we did not determine whether differences in neuronal activity drive the OLLC phenotype observed here. Future studies directly manipulating glutamatergic signaling and neuronal activity will be important for determining whether altered OLLC communication with ExNs contributes to the persistence of OLLC_3-SID state, and whether normalization of these interactions can promote productive myelination. In addition, our study did not distinguish the relative contributions of trisomic OLLC-specific cell-intrinsic differences from those originating in the surrounding neuronal environment, which should be investigated in the future studies.

Together, our findings identify a previously unrecognized divergence in human oligodendrocyte development in DS, characterized by enrichment of a differentiated, neuron-interacting pre-myelinating state without corresponding activation of myelination-supportive programs. The correspondence of this state with transcriptional programs in the developing human DS cortex, together with altered neuron–OLLC interactions and myelin ultrastructure in FAs, supports a model in which disrupted coordination between oligodendrocyte differentiation and neuronal engagement may contribute to impaired myelination in DS.

## Methods

### iPSC lines and maintenance

For these studies, we used three isogenic DS-derived paired induced pluripotent stem cell (iPSC) lines. We cultured two female paired cell lines (euploid WC-24-02-DS-B and trisomic WC-24-02-DS-M and euploid ILD11-3 and trisomic ILD1-2) previously used in our laboratory^33,145^ and one male paired cell line (euploid DS1-iPS4-disomic and DS1-iPS4-trisomic). The WC-24-02-DS-B and WC-24-02-DS-M cell lines were generated by Dr. Anita Bhattacharyya’s laboratory at the University of Wisconsin, while ILD11-3 and ILD1-2 cell lines were generated by Dr. Joaquin M Espinosa’s laboratory at the University of Colorado Anschutz Medical Campus, and both isogenic pairs were generously shared with our lab. The DS1-iPS4-disomic and DS1-iPS4-trisomic cell lines were generated by Dr. Orkin’s laboratory^146,147^ and acquired through Boston Children Hospital. All euploid cell lines exhibit a normal karyotype and all trisomic lines exhibit a normal karyotype aside from the triplicated Hsa21. Each pair of isogenic cell lines was used for three independent FA differentiation experiments. iPSCs were cultured on Matrigel-coated plates (cat. 354277 Corning, Corning, NY, USA), replacing mTeSR Plus media (cat. 100-0276, STEMCELL Technologies, Vancouver, BC, Canada) every other day. iPSCs were passaged with ReLeSR (cat. 100-0483, STEMCELL Technologies) and only iPSCs with between 24 to 50 passages that displayed normal morphology and tested negative for mycoplasma were used for the organoid generation.

### Generation of dorsal and ventral cortical organoids (dCOs, vCOs)

<u>dCO generation</u>: dCOs were generated as previously described^29,33,34^. Briefly, one well of iPSCs from a 6-well plate was rinsed with 1 mL phosphate-buffered saline (PBS) and dissociated by incubation with 1 mL Accutase (cat. 7920, STEMCELL Technologies) for approximately 5 min. Cells were gently triturated to obtain a single-cell suspension, diluted with 5 mL DMEM/F12 (cat. 11330–057, ThermoFisher Scientific, Waltham, MA, USA), centrifuged for 5 min, and resuspended in 1 mL mTeSR Plus supplemented with 50 µM ROCK inhibitor (cat. 12-541-0, Fisher Scientific, Pittsburgh, PA, USA) for cell counting. Cells were then plated into low-adhesion V-bottom 96-well plates (cat. MS-9096VZ, S-Bio Prime, Constantine, MI, USA) at a density of 15,000 cells per well in 150 µL of medium.

At 1 *DIV* (days *in vitro)*, medium was replaced with 100 µL TeSR-E6 (cat. 0596, STEMCELL Technologies) supplemented with 2.5 µM dorsomorphin (DM; cat. P5499, MilliporeSigma, Burlington, MA, USA) and 10 µM SB-431542 (SB; cat. S4317, MilliporeSigma). Media containing DM and SB was replaced daily between 3 and 6 *DIV*.

At 7 *DIV*, 100 µL of medium was replaced with Neural Medium (NM) consisting of Neurobasal-A (cat. 10888022, Life Technologies, Carlsbad, CA, USA) supplemented with B-27 without vitamin A (1:50; cat. 12587, Life Technologies), GlutaMax (1:100; cat. 35050061, Life Technologies), penicillin/streptomycin (100 U/mL; 1:100; cat. 15140122, Life Technologies), and Primocin (1:500; cat. ant-pm-1, InvivoGen, San Diego, CA, USA). From 7 to 25 *DIV*, NM was supplemented with fibroblast growth factor-2 (FGF2; 20 ng/mL; cat. 233-FB-25/CF, R&D Systems, Minneapolis, MN, USA) and epidermal growth factor (EGF; 20 ng/mL; cat. 236-EG-200, R&D Systems). Between 16 and 24 *DIV*, cortical organoids (COs) were transferred to 24-well ultra-low attachment plates (cat. 07-200-602, ThermoFisher Scientific) and cultured individually to prevent the fusion of the organoids. From 25 to 39 *DIV*, NM was supplemented with brain-derived neurotrophic factor (BDNF; 20 ng/mL; cat. AF-450-02, PeproTech, Waltham, MA, USA), neurotrophin-3 (NT3; 20 ng/mL; cat. 450-03, PeproTech), and 1% Geltrex (cat. A1569601, Life Technologies). Half-medium changes were performed every other day. Between 40 and 50 *DIV*, dCOs were maintained in NM containing 1% Geltrex, with half-medium changes every two days. To support the population of oligodendrocytes within dCOs, platelet-derived growth factor AA (PDGF-AA; 10 ng/mL, cat. 221-AA, R&D Systems) and insulin-like growth factor (IGF; 10 ng/mL, cat. USA291-GF-200, R&D Systems) were added to NM media between 50 to 60 *DIV*. 3,3′,5-triiodothronine (T3; 40 ng/mL, cat. T6397, MilliporeSigma) was added to the NM between 61 to 69 *DIV*, and media changes were performed every two days. From 70 *DIV* onwards, the dCOs were cultured in NM with the media changes performed every two days. Dorsal identity of the dCOs was validated by immunohistochemistry at day 50 through detection of PAX6 and lack of NKX2.1 expression.

#### vCO generation

vCOs were generated as previously described^28,43,148^ by introducing ventral patterning cues into the dCO differentiation protocol. vCO differentiation followed the same procedures as dCOs from 0 to 6 *DIV*, with the addition of the Wnt pathway inhibitor IWP-2 (5 µM; cat. S7085, Selleckchem, Houston, TX, USA) from 4 to 24 *DIV*. Beginning at 7 *DIV*, vCOs were transitioned to differentiation and maintenance medium (DMM) containing DMEM/F12 supplemented with B-27 without vitamin A (1:50), N2 supplement (cat. 17502048, ThermoFisher Scientific), human insulin (25 µg/mL; cat. I9278-5ML, MilliporeSigma), Minimum Essential Medium Non-Essential Amino Acids Solution (MEM NEAA; 1:100; cat. 11140076, ThermoFisher Scientific), penicillin/streptomycin (1:100), GlutaMax (1:100), and β-mercaptoethanol (0.1 mM; cat. M3148, MilliporeSigma).

From 7 to 24 *DIV*, DMM was supplemented with EGF (20 ng/mL) and FGF2 (20 ng/mL). To induce ventral identity, the Sonic Hedgehog (SHH) pathway smoothened agonist (SAG; 1 µM; cat. 566660, MilliporeSigma) was added between 12 and 24 *DIV*. Neuronal expansion and oligodendrocyte enrichment were promoted between 24 and 37 *DIV* by supplementing DMM with BDNF (20 ng/mL), NT3 (20 ng/mL), PDGF-AA (10 ng/mL), IGF (10 ng/mL), T3 (60 ng/mL), hepatocyte growth factor (HGF; 5 ng/mL; cat. 315-23, PeproTech), cyclic AMP (cAMP; 1 µM; cat. D0627, MilliporeSigma), and biotin (100 ng/mL; cat. B4639, MilliporeSigma).

From 37 *DIV* onward, vCOs were maintained in DMM supplemented with T3, cAMP, biotin, and ascorbic acid (AA; 20 µg/mL; cat. A4403, MilliporeSigma), with media changes performed twice weekly. Ventral identity of the vCOs was validated by immunohistochemistry at day 50, through detection of NKX2.1 and absence of PAX6 expression.

Following the original protocols, organoid cultures were maintained under static conditions, in the incubator at 37°C supplemented with 5% CO_2_.

### Forebrain assembloid (FA) generation

At 75 *DIV*, one vCO and one dCO from the same cell line and differentiation experiment were fused as previously described^43^. COs were placed in close contact to encourage fusion and co-cultured for 4 days to allow physical adhesion. If complete fusion was not achieved after 4 days, the co-incubation period was extended for up to 8 days to allow stable fusion between dCOs and vCOs. Media was changed in tubes every other day until fusion. Once fused, FAs were transferred back to 24-well ultra-low attachment plates and were maintained with half-volume media changes every other day^149^.

### Viral transduction of vCOs and FAs

*vCOs* were transduced with pAAV-Sox10-tdTomato (titer ≥ 1×10^13^ vg/mL, cat. AAV1S(VB230712-1610aft)-K, VectorBuilder, Chicago, IL, USA)^44^ at 60 *DIV*, prior to fusion with the dCOs. For the transductions, vCOs were incubated individually with ∼20 µL media and 0.5 µL of virus in 1.5 mL Eppendorf tubes for 40 min, followed by the addition of 100 µL media for overnight incubation^46^. The following day, the vCOs were transferred back to 24-well ultra-low attachment plates with 500 µL media.

### Cryoprotection and slice preparation

dCOs, vCOs, and FAs were fixed overnight in 4% paraformaldehyde (PFA) at 4 °C, washed 3 times with PBS, then transferred to 30% sucrose solution. dCOs, vCOs, and FAs were embedded in mixture of 30% sucrose and Tissue-Plus O.C.T. Compound (OCT: optimal cutting temperature medium; cat. 23-730-571, Fisher Scientific) at a 40:60 ratio. dCOs, vCOs, and FAs were flash frozen, stored at -80°C, and sectioned at 12 µm or 300 µm.

### Immunohistochemistry (IHC)

On slide: Slides with dCOs, vCOs, and FAs were rehydrated and washed 3 times for 10 minutes each with PBS with 0.25% Triton X-100 (PBST; cat. BP151-500, Fisher Scientific). The slides were incubated in blocking solution with 3% donkey serum (cat. D9663, Sigma Aldrich) in PBST for 1 hour. Antigen retrieval was performed by microwaving slides in blocking solution twice at 25 °C for 5 minutes at 250W in a PELCO Biowave Pro+ microwave (cat. 36700, Ted Pella, Inc., Redding, CA, USA), allowing slides to cool to RT in between microwave cycles. Slides were then incubated overnight with primary antibodies (**Supplementary Table 12**) diluted in blocking solution at 4 °C. Slides were washed 3 times for 10 minutes each with PBST, incubated with secondary antibodies (**Supplementary Table 12**) diluted in blocking solution for 1 hour at RT, washed 3 times for 10 minutes each, and coverslipped with ProLong Gold Antifade Mountant (cat. P36930, ThermoFisher Scientific) or ProLong Gold Antifade Mountant with DAPI (cat. P36931, ThermoFisher Scientific).

Free floating: 300 µm sections with FAs were washed 3 times for 10 minutes each with 0.01M PBS and incubated for 1 hour in 50 mM glycine in 0.01M PBS. Sections were washed 3 times for 10 minutes each with 0.01M PBS, antigen retrieval was performed with sodium citrate (pH = 6) for 20 minutes at 55-65 °C, and then cooled for 30 minutes. Sections were washed 3 times for 10 minutes each, then blocking was performed with 5%BSA-NDS for 1 hour. Primary antibodies (**Supplementary Table 12**) were diluted in BSA-c/NDS incubation buffer-TxSIn and incubated for 72 hours with 2 microwave sessions at 150W and 40 °C for 10 minutes during the incubation. Sections were washed 3 times for 10 minutes each and secondary antibodies (**Supplementary Table 12**) diluted in BSA-c/NDS incubation buffer-TxSIn were incubated overnight, microwaved twice, washed again 3 times for 10 minutes each, and coverslipped with ProLong Gold Antifade Mountant with DAPI (cat. P36931, ThermoFisher Scientific).

### Imaging and quantification Whole organoid analysis

Z-stacks with a step size of 1 µm from at least 3 sections for each dCO, vCO, or FA were obtained using Slideview VS200 slide scanner (Evident Scientific, Waltham, MA, USA) with 20x objective. Whole section maximum intensity projections (MIPs) were analyzed using QuPath v0.7.0 (Bankhead, Loughrey et al. 2017). For each image, total cell number was counted based on DAPI expression using the “Cell detection” function. Next, the threshold for each viral or IHC label was manually set using the “Create single measurement classifier” function using “nucleus: mean” for nuclear markers and “cell: mean” for cytoplasmic markers. Once all markers were classified, object classifiers were applied sequentially to detect positivity and colocalization of markers. Measurements were documented with the “Show annotation measurements” function. Counts from the 3 sections of the same CO or FA were averaged and reported as a proportion of total cells per CO or FA.

### Confocal imaging

High-resolution confocal z-stack tile scans (1 µm optical sections) were acquired using an Andor BC43 confocal microscope (Oxford Instruments, Belfast, UK) with a 40× Plan Fluorite oil immersion 1.3 NA, 0.24mm WD objective. Images were deconvolved and stitched using Fusion Benchtop software (Oxford Instruments) and displayed as maximum-intensity projections (MIPs).

### Synaptic marker analysis

Z-stacks with a z-step size of 0.1 µm from 3 separate fields of view (FOVs) (over 2-3 sections) per FA were obtained with an Andor BC43 confocal microscope (Oxford Instruments, Belfast, UK) with a 40x Plan Fluorite oil immersion 1.3 NA, 0.24mm WD objective, and deconvolved using Fusion Benchtop software. Volumetric renderings of oligodendrocytes and their synaptic elements were generated by creating ‘Surface’ representations for BCAS1, VGLUT1, GluA4, HOMER1b/c and CC1, and ‘Spots’ representations for PSD95 and VGAT in Imaris v10.2.0 (Bitplane, Belfast, UK). Spots were filtered with the ‘quality’ parameter. Imaris machine learning segmentation was then used to define spots and surfaces.

Contacts were defined as spots (VGAT) or surfaces (VGLUT1) within 1 µm of the BCAS1 or CC1 surface, and were normalized to BCAS1, CC1, or GluA4-BCAS1 surface area. Surface-surface colocalizations (GluA4 and HOMER1b/c) were defined as overlapped volumes with BCAS1 > 0.01 µm^3^ and new surfaces were created for the overlapping volumes (GluA4-BCAS1 and HOMER1/bc-BCAS1), which were normalized to BCAS1 volume. Spot-surface colocalization (PSD95) was defined as spots within 0.01 µm of the BCAS1 surface and were normalized to BCAS1 surface area (PSD95-BCAS1). Image reconstructions were created using ImageJ/FIJI for background subtraction and smoothing with Median filtering, and Imaris v10.2.0 (Bitplane) for 3D ‘Surface’ representations and rotations.

### Migration

#### Live-cell image acquisition

At 55 *DIV*, vCOs were transduced prior to FA fusion with pAAV-Sox10-tdTomato (titer ≥1 × 10^13^ vg/mL, cat. AAV1S(VB230712-1610aft)-K, VectorBuilder, Chicago, IL, USA) as previously described^44^. At 120 *DIV*, FAs containing *SOX10-tdTomato* were transferred to glass bottom 96 well plates (cat. P96-1.5H-N, Cellvis, Mountain View, CA, USA). At 120 *DIV*, fluorescent live-assembloid imaging was performed on a Nikon CSU-W1 SoRa spinning disc confocal microscope (Nikon, Tokyo, Japan) equipped with a S PLAN FLUOR LWD 20× objective (0.7 NA, 2.3 mm working distance), controlled using NIS-Elements software. FAs were maintained in an environmental chamber at 37 °C and 5% CO_2_. For time-lapse recordings, we acquired 20 µm z-stacks every 20-30 minutes over a 17 - 18-hour imaging period, resulting in 4D images. For multitile acquisitions, adjacent fields were acquired with 15% overlap and stitched using NIS Elements built-in method.

### 4D image pre-processing

A pipeline recently developed in our laboratory was used for all live-cell image processing^43^. All image pre-processing was performed using custom MATLAB scripts with ImageJ used for visualization. Raw image data were organized as multi-dimensional hyperstacks containing spatial (x, y, z) and temporal (t) dimensions, and channel indices. To correct for uneven illumination, background gradients, and low-frequency intensity variations, a 3D Gaussian blur was applied to estimate background signal. For each time point, the image volume was convolved with a 3D Gaussian kernel (σx = 21 pixels, σy = 21 pixels, σz = 9 pixels), corresponding to an effective smoothing scale of approximately 13.65 µm laterally and 180 µm axially based on the imaging resolution. The resulting smoothed volume captured slowly varying background signal while preserving the spatial scale of cellular features. The blurred volume was subtracted from the original data, enhancing contrast and improving visibility of individual labeled calls across dense assembloid regions, while suppressing gradual background variations. Bulk sample drift was then corrected in FIJI/ImageJ using the Correct 3D Drift plugin^150^, applying automated rigid registration in X, Y and Z relative to the first timepoint.

### Manual cell tracking

Manual cell tracking was performed in ImageJ/FIJI using the OrthoTrack plugin^151^, which enables precise annotation of cell positions across three spatial dimensions and time. Pre-processed image stacks were converted to 4D hyperstacks (x, y, z, t) and individual cells were manually tracked by identifying the cell center in orthogonal views (XY, XZ, and YZ) at each time point. Cells were included in the analysis if they were detected in all time frames throughout the imaging period, were located near the vCO-dCO border and exhibited movement directionality from the vCO toward dCO as previous described^43^. For each tracked cell, 3D coordinates were recorded at every time point for quantitative analysis. Each assembloid was tracked by at least 3 individuals and a minimum of 3 assembloids per genotype per cell line were analyzed.

### Analysis of tracked cells

Migration metrics were calculated from manually tracked cell coordinates using a custom MATLAB analysis pipeline. For each tracked cell, 3D trajectories were reconstructed using x, y, and z positions across time. Total distance traveled was calculated as the cumulative sum of Euclidean distances between consecutive time points. Relative displacement was computed as the straight-line distance between the initial and final cell positions as a measure of net migration. Cell migration speed was calculated by dividing frame-to-frame displacement by the imaging interval, and average speed was computed over the full imaging duration.

### Transmission electron microscopy (TEM)

At 150 *DIV*, FAs were collected, washed with PBS 2 times and fixed in 4% PFA with 0.2% glutaraldehyde in 0.1M Cacodylate buffer for at least 24 hours. FAs were cut in half and rinsed with cacodylate buffer 3 times over 1-2 hours. Samples were incubated in 1.5% KFe:1% osmium solution in 0.1M PB (phosphate buffer) on rotator for 30 minutes, then rinsed with 4 times with 0.1M PB (phosphate buffer). Samples were then incubated in 1% osmium in 0.1M PB on rotator for 30 minute, rinsed 4 times in 0.1M PB, then 3 times with distilled water. Dehydration was performed with 2 washes of 50% EtOH for 5 minutes each, incubation in 2% uranyl acetate in 70% EtOH for 40 minutes, 2 washes of 85% EtOH for 5 minutes each, 2 washes of 95% EtOH for 5 minutes each, and 3 washes of 100% EtOH for 10 minutes each. Samples were infiltrated on rotator overnight in 1:1 EM812 resin to propylene oxide then transferred to 100% resin with DMP-30 for 4 hours in vacuum desiccator. Finally, samples were embedded in 100% resin and cured at 60°C for 72 hours. Ultrathin 70nm sections were cut using an Ultracut UCT ultramicrotome (Leica, Wetzlar, Germany) and collected on Gilder fine bar 200 mesh copper grids (cat. G200HS, Ted Pella). Sections were visualized on a JEOL JEM-1400 Flash transmission electron microscope (JEOL, Tokyo, Japan). For each sample, one grid square equaling 12769 µm^2^ (113 µm × 113 µm) was examined and all notable features were imaged.

TEM images were analyzed in ImageJ for both quantitative and qualitative metrics. Myelin was classified based on level of compaction (loose/early, mid-compact, and compact). Axon area, diameter, and myelin thickness were measured using the FIJI polygon tool. Thickness measurements were analyzed using k means to detect subpopulations within our data. Intracellular and extracellular axons were determined qualitatively using surrounding structures such as mitochondria, nuclei, and cytoplasm to determine location. Defective myelin sheaths were identified by presence of either detachment of myelin sheaths from the axon, dark axoplasm, irregular lamellar splitting or redundant myelin formation.

### Statistical analysis

Quantitative data are expressed as mean ± SEM. Statistical analyses of IHC and live cell migration data were performed with R version 4.5.2 in RStudio and visualized in GraphPad Prism 10.6, and values of p < 0.05 were considered significant. To account for the nested structure of our data (3 isogenic paired cells lines, 2-3 independent differentiations per line and multiple FAs per differentiation), we used a linear mixed-effects model (LME) implemented with the *lme4* and *lmerTest* packages with genotype as a fixed effect. Cell line and differentiation were included as random effects. The primary model was specified as:

Outcome ~ Genotype + (1 | Cell Line) + (1 | Cell Line:Differentiation)

COs or FAs were biological replicates, Cell Line represented independent iPSC genetic backgrounds and Differentiation referred to independent differentiation experiments nested within each cell line. Fixed-effect significance was assessed using t-tests with degrees of freedom estimated by the *lmerTest* package. The contributions of genotype, cell line, and differentiation to model fit were additionally evaluated using likelihood ratio tests. To determine whether genotype effects differed across genetic backgrounds, models including Genotype × Cell Line interaction were compared with models lacking the interaction. Effects of genotype within individual cell lines were further analyzed using estimated marginal means and pairwise contrasts with the *emmeans* package. Variance components were extracted from the fitted mixed-effects models to assess the proportion of variability attributable to cell line, differentiation, and residual variation.

TEM data were analyzed using GraphPad Prism 10.6. Normality of data was analyzed using Shapiro-Wilk test, and normally distributed datasets were compared using unpaired t-tests with Welch’s correction while non-normally distributed datasets were compared using Mann Whitney tests. Outliers were removed where appropriate using ROUT (Q=1%) and values of p < 0.05 were considered significant.

### RNA sequencing

#### Cell dissociation for RNA sequencing

Cell dissociation was performed using the Papain Dissociation System (cat. LK003150, Worthington Biochemical Corporation, Lakewood, NJ, USA), as previously performed in our laboratory^34,48^. Four FAs from 2 independent differentiations (148 *DIV* and 151 *DIV*, hereafter referred to as 150 *DIV*) were transferred to 1 well of an ultra-low attachment plate (cat. 07-200-601, Corning) with 500 µL Neurobasal A media and cut into small pieces. During a 40-minute incubation period in papain solution, samples were pipetted 7 times with a wide bore P1000 and 2 times with a regular P1000 every 10 minutes. Then, the mixture was triturated vigorously with a P1000 pipette and the well was washed with an additional 200 µL of papain solution. The solution was strained through a 40-µm strainer (cat. 431750, Corning) and centrifuged at 300g for 2 min at RT. The supernatant was aspirated and cells were resuspended in 2 mL in combination of Earle’s Balanced Salt Solution (EBSS), reconstituted albumin-ovomucoid inhibitor solution and DNase solution. The mixture was centrifuged at 250 g for 3 min at RT, resuspended in 1 mL 1% BSA in PBS, centrifuged again at 250 g for 3 min at RT, and resuspended in a final mixture of 1 mL 0.04% BSA in PBS. Cells were pipetted 10 times, counted on Countess II FL Automated Cell Counter (AMQAF1000, ThermoFisher Scientific) using Trypan blue stain (cat. T10282, ThermoFisher Scientific). Only samples with greater than 85% viability were used for subsequent steps. 20% of each sample was used for single-cell RNA sequencing (scRNA-seq), with the target capture of 4000 cells per sample. The remaining 80% of each sample was processed for nuclei isolation using the Chromium Next GEM Single Cell Multiome ATAC + Gene Expression Sequencing kit (10x Genomics, Pleasanton, CA, USA).

FAs from each isogenic cell line were profiled by both scRNA-seq and snRNA-seq, which capture complementary aspects of cellular transcriptomes. scRNA-seq captures both cytoplasmic and nuclear transcripts and typically provides higher detection depth^152,153^, while snRNA-seq captures predominantly nuclear, often intronic, transcripts at lower depth^154^. Each cell-line pair was analyzed separately within each technology, yielding six matched euploid–trisomic comparisons across three genetic backgrounds. Downstream analyses were performed within these matched pairs and integrated across them, so that reported genotype- and state-associated differences were those reproducible across both technologies and all three genetic backgrounds rather than specific to one technology^153^.

### Preparation of samples for single-cell RNA sequencing (scRNA-seq)

cDNA synthesis, amplification and library preparation for scRNA-seq from dissociated cells was performed by the Boston University Chobanian & Avedisian School of Medicine, Microarray and Sequencing Resource (MSR) Core using 10x Genomics Chromium Single Cell 3′ Library and Bead Kit (v3; 10x Genomics) according to manufacturer’s instructions.

### Preparation of samples for single nucleus RNA sequencing (snRNA-seq)

Nuclei isolation protocol was adapted from 10x Nuclei Isolation for Single Cell Multiome ATAC + Gene Expression Sequencing (10x Genomics; CG000365 Rev C). 100,000 to 1,000,000 cells were added to a 1.5 mL microcentrifuge tube and centrifuged at 300g for 5 minutes at RT. Supernatant was removed, cells were resuspended in 100 µL chilled 0.1% Lysis Buffer (10x Genomics), and pipette mixed 10 times. Samples were incubated on ice for 3 minutes, and 500 µL chilled Wash Buffer (10x Genomics) was added to lysed cells before pipette mixing 3-5 times. Lysed cells were centrifuged at 500 g for 5 minutes at RT. Process was repeated for a total of 3 washes. Nuclei were resuspended in 14-20 µL Diluted Nuclei Buffer (10x Genomics), and nuclei concentration and integrity were determined by manual counting with a hemocytometer by BUMSR Core. Samples with nuclei integrity above 80% were used for subsequent processing, with a target capture of 4000 nuclei per sample.

Any subsequent sample preparation for sequencing followed 10x Chromium Next GEM Single Cell Multiome ATAC + Gene Expression protocol (10x Genomics; CG000338 Rev F) including Transposition, GEM Generation and Barcoding, Post GEM Incubation Cleanup, Pre-Amplification PCR, ATAC Library Construction, cDNA Amplification, Gene Expression Library Construction.

Sequencing was performed by BUMSR core using NextSeq 2000 with P4 Flow Cell (Illumina; San Diego, CA, USA) according to manufacturer’s instructions.

### Bioinformatic analysis Quality control

scRNA-seq and snRNA-seq data from FA samples at 150 *DIV* were processed in R using the Single Cell Toolkit (SCTK)^155^. Cells and nuclei with mitochondrial gene content greater 10% were removed. Putative doublets were identified with scDblFinder and removed. DecontX was used to estimate ambient RNA^156^. Cells and nuclei with contamination scores greater than 80% were excluded.

#### Integration, Clustering, and cell type annotation

For integration of scRNA-seq and snRNA-seq data, we combined the corresponding scRNA-seq and snRNA-seq Seurat objects (Seurat V 5.4.0) into a single merged object. These datasets comprised three paired isogenic cell lines (WC, ILD, DS1), each represented in both technologies and in euploid and trisomic genotypes. We applied SCTransform normalization^157^ followed by principal component analysis (PCA) with 30 components, and then used Harmony for batch correction and dataset integration^158^. The sequencing technology (scRNA-seq versus snRNA-seq) was specified as the grouping variable and using a reduced θ parameter (θ = 3) to limit overcorrection while aligning shared biological structure between technologies. The Harmony-corrected embedding was used as the input for UMAP (30 Harmony dimensions) to obtain a joint low-dimensional representation of all cells, and the same Harmony dimensions were used to construct a shared nearest-neighbor graph, on which Seurat’s graph-based clustering was performed^159^. Clusters were identified at a resolution of 0.4, yielding stable, biologically interpretable cell types across the integrated scRNA-seq and snRNA-seq data, and UMAPs colored by dataset, cell line, genotype, and cluster identity were inspected to confirm adequate mixing of technologies and consistent separation of major cell populations.

Cell types in the integrated scRNA-seq/snucRNA-seq dataset were annotated using well-established marker genes for major neural and glial populations and their developmental states as indicated in **Supplementary Table 2**. For excitatory neurons, we used genes associated with cortical identity and layer specialization such as *NEUROD2, NEUROD6, SNAP25, SATB2, CUX2, RORB, BCL11B, FEZF2*. Progenitor and radial glial populations were identified by markers including *EOMES, VIM, SOX2, HES1, MKI67* and *TOP2A*, while astrocytes were defined by expression of *SPARCL1, S100B, AQP4* and *GFAP*, and OLLCs by *OLIG2, PDGFRA,* and *ITPR2.* Inhibitory interneurons were annotated based on the expression of *GAD1, GAD2, DLX2,* while Reelin+ cells and choroid plexus (ChP) cells were identified based on the expression of *RELN* and *TTR,* respectively.

### Cell-type composition analysis

To test for genotype-associated differences in cell-type composition while accounting for sample-level replication, per-sample cell-type proportions were compared between euploid and trisomic FAs using propeller (speckle R package)^160^. Each cell line × sequencing technology combination was treated as an isogenic pair, and a paired design (∼ 0 + genotype + pair) was used. P values were corrected across cell types using the Benjamini–Hochberg method, and cell types with FDR < 0.05 were considered significantly different.

### Entire transcriptomic object alignment with human fetal cortex reference single-cell dataset

To assess how closely FA cell types resembled their *in-vivo* counterparts, we compared them with a human fetal cortex reference dataset with both DS and control samples^52^. After log-normalization, the 2,000 most variable genes were identified in the fetal reference (Seurat FindVariableFeatures, vst method) and matched to the genes in the FA dataset. We calculated average expression for each fetal cell cluster, as defined and annotated by the original authors to represent distinct cell types, and for each FA cell type, with genotypes pooled. We then computed Spearman correlations between every fetal cluster and every FA cell type. Because correlations were based on cluster-level averages, they reflect relative similarity between cell types rather than single-cell correspondence, and each assembloid cell type was interpreted relative to its best-matching fetal populations rather than by absolute correlation values.

### OLLC subclustering

During subclustering of the glial subset (clusters 4 and 5), we initially obtained 11 glial clusters (0–10), using resolution 0.4. One small cluster (cluster 10) also exhibited high expression of astrocyte markers and low expression of OLLC markers. We removed the cells belonging to cluster 10 and repeated the subclustering so that downstream analyses could be performed on a clearer separation of these two glial lineages. This dataset contained 1,985 cells from three cell lines, including 905 euploid and 1,080 trisomic cells profiled by scRNA-seq and snRNA-seq respectively. Because the OLLC subset was relatively small and showed uneven representation across cell lines and sequencing technologies, we performed variable-gene selection and integration specifically within this population. Cells were first separated into six groups based on cell line and assay type: WC single-cell, WC single-nucleus, ILD single-cell, ILD single-nucleus, DS1 single-cell, and DS1 single-nucleus. To reduce the separation driven by technical differences and individual lines, within each group, the top 5,000 highly variable genes (HVGs) were identified. The intersection of HVGs across all six groups resulted in 1,155 genes which was used for clustering in the Seurat workflow (**Supplementary Table 6**). Harmony integration was applied to the first 15 principal components using cell line and dataset as integration variables, and the first 15 Harmony dimensions were used for UMAP visualization, nearest-neighbor graph construction and graph-based clustering at a resolution of 0.4.

### Cluster Marker Analysis

Cluster-specific marker genes were identified for the Harmony-derived OLLC subclusters using Seurat FindAllMarkers with the Wilcoxon rank-sum test. Marker testing was restricted to the 1,155 highly variable genes shared across the six cell line and sequencing-technology groups, thereby focusing the analysis on genes with reproducible variability across the dataset. Only positively enriched markers detected in at least 25% of cells in either the target cluster or the comparison cells were tested (https://satijalab.org/seurat/reference/findallmarkers).

For each cluster, markers were further filtered to retain genes with an average log_2_ fold change greater than 0.25 and expression in fewer than 50% of cells outside the cluster (pct.2 < 0.5). This additional filtering was used to prioritize genes with relatively specific expression patterns rather than genes broadly expressed across multiple OLLC states. Ensembl gene identifiers were mapped to gene symbols using org.Hs.eg.db; where a gene symbol was unavailable, the original Ensembl identifier was retained. Nominal Wilcoxon test *p* values were adjusted using the Benjamini-Hochberg method to calculate false discovery rates. The 10 highest-ranking specific markers per cluster, ordered by average log_2_ fold change, were used for visualization in a heatmap of scaled expression values.

To identify biological processes associated with each OLLC cluster, Gene Ontology Biological Process gene set enrichment analysis (GSEA) was performed using gseGO in clusterProfiler. For each cluster, a one-versus-rest comparison was generated using Seurat FindMarkers across the same shared highly variable gene set. Genes were ranked by average log_2_ fold change, including both positively and negatively associated genes, and duplicate gene symbols were removed prior to enrichment analysis. Gene Ontology Biological Process terms containing 5 to 500 genes were tested using 10,000 permutations and Benjamini-Hochberg correction. Only positively enriched pathways with FDR-adjusted *p* < 0.05 were retained. GSEA tests whether genes within predefined pathways are non-randomly concentrated toward one end of a ranked gene list^161^. The 10 most significantly enriched terms per cluster were displayed in a lollipop plot, with normalized enrichment score indicating the magnitude and direction of enrichment and point size representing adjusted significance.

### Genotype Enrichment Analysis

To assess genotype-biased representation across OLLC subclusters, we quantified the proportion of trisomic and euploid cells within each subcluster and tested for significant enrichment using Fisher’s exact test. For each subcluster, a 2×2 contingency table was constructed comparing the number of cells of each genotype in that subcluster versus all remaining subclusters. A pseudocount of 0.5 was added to cells where any count was zero to avoid undefined odds ratios. P-values were corrected for multiple comparisons using the Benjamini–Hochberg (BH) false discovery rate method. Effect sizes were quantified as log_2_ fold-change (log_2_FC) of genotype proportions, Cohen’s h, and log_2_ odds ratio. Subclusters were considered significantly enriched if FDR < 0.05 and |log_2_FC| ≥ 0.5 or |Cohen’s h| ≥ 0.3. Results were visualized as a volcano plot with log_2_FC on the x-axis and −log_10_(FDR) on the y-axis, with dashed lines indicating effect size (|log_2_FC| = 0.5) and significance (FDR = 0.05) thresholds. All analyses were performed in R using the *Seurat*, *dplyr*, *ggplot2*, and *ggrepel* packages.

### Differential Gene Expression (DEG) Analysis

Differential expression between OLLC_2 and OLLC_3 was assessed using scMetaIntegrator, a paired single-cell meta-analysis framework. OLLC_2 was enriched for euploid cells, whereas OLLC_3 was enriched for trisomic cells. Because OLLC subclusters also differed in their contribution from each cell line, each cell line and sequencing-technology combination was treated as a paired comparison group. Within each group, cells from OLLC_2 and OLLC_3 were assigned separate cluster-specific sample identifiers, and only groups containing both clusters were included. Gene-level effect sizes (Hedges’ g) were calculated within each pair and combined across pairs, and p-values for increased and decreased expression were combined using Fisher’s method and corrected using the Benjamini–Hochberg method. This approach accounts for cell-level variation while avoiding the treatment of individual cells as independent biological replicates. Positive effect sizes indicate higher expression in OLLC_3 and negative effect sizes higher expression in OLLC_2. Genes tested in all six pairs with a combined Fisher FDR < 0.05 in either direction were considered differentially expressed. To identify genotype differences across the whole OLLC population, the same scMetaIntegrator approach was applied to all OLLCs, comparing trisomic with euploid cells within each pair. GO Biological Process over-representation analysis was performed separately on genes higher and lower in trisomic OLLCs using enrichGO in clusterProfiler, with all genes tested in all six pairs as the background and BH-adjusted p < 0.05. The 40 most significant terms in each direction were displayed as networks, in which nodes represent GO terms (size = number of DEGs) and edges connect terms sharing genes (Jaccard similarity ≥ 0.2).

#### Gene Set Enrichment Analysis (GSEA)-based Gene Ontology (GO) enrichment Analysis

For GSEA, genes were ranked by average log2 fold change between OLLC_3 and OLLC_2 using a pooled Seurat Wilcoxon rank-sum test. These statistics were used exclusively for gene ranking, while DEGs were defined by scMetaIntegrator as described above. GSEA was performed with gseGO in clusterProfiler (https://yulab-smu.top/biomedical-knowledge-mining-book/021-go.html) to identify enriched GO Biological Process (BP) terms, with significance determined using Benjamini–Hochberg-adjusted FDR to identify significant terms. Positive normalized enrichment scores (NES) indicated enrichment in OLLC_3, whereas negative NES indicated enrichment in OLLC_2. NES accounts for differences in gene-set size and allows enrichment strength to be compared across pathways^162^. Significant BP terms were grouped into broader functional categories based on shared biological annotation. For visualization, the 50 terms with the highest absolute NES were displayed separately for each OLLC state, ranked by enrichment magnitude and colored according to functional category.

### Celda Module Analysis

To examine transcriptional heterogeneity within the refined OLLC population, we applied the celda_CG co-clustering model from Celda^163^ (https://bioconductor.org/packages/celda/). Co-expression structure was inferred from count data using minimum thresholds of 5 cells per gene and 5 counts per feature, considering up to 5,000 genes and specifying 30 cell states and 40 gene modules, with genotype included as a sample-level annotation during model fitting. Celda-derived cell groupings and gene modules were visualized in the existing Seurat UMAP embedding (https://satijalab.org/seurat/), and representative genes for each module were identified from feature rankings for downstream interpretation. To assess known functional relationships among genes within Celda modules M13 and M2, we built gene association networks using the STRING database^164^ (version 12.0) through the STRINGdb R package (https://bioconductor.org/packages/STRINGdb). Module genes were mapped to their corresponding human STRING entries (taxonomy ID 9606). Associations between genes within each module were retained if their combined score was ≥ 400 (medium confidence).

### Module Enrichment in OLLC Subclusters

To summarize Celda-derived transcriptional modules across OLLC subclusters, we performed a module enrichment analysis at the cluster level using Celda^163^. For each module, we computed per-cell z-scored expression and used Wilcoxon tests to identify modules that were significantly over- or under-expressed in each OLLC cluster, retaining the top modules per cluster based on FDR-adjusted *p* values. These modules were then subjected to GO-based functional annotation using GO and KEGG terms were initially screened using DAVID-like criteria (*p* < 0.1, at least five overlapping genes), and terms with FDR-adjusted *p* < 0.05 were retained for downstream annotation. This was followed by manually grouping terms into broader functional categories (**Supplementary Tables 10, 11**). M7 was removed from the analysis as it was predominantly composed of uncharacterized transcripts, limiting its functional interpretation and M37, which was only modestly enriched in OLLC_2-TMS and was characterized by broad transcriptional and chromatin-remodeling programs, was not assigned to a higher-order categories. Alluvial plots were generated for the OLLC clusters of interest to show which Celda modules mapped to each functional category and which clusters most strongly expressed those modules.

### Slingshot Pseudotime

For pseudotime analysis, we focused on the OLLC object containing 6 clusters (OLLC_0– OLLC_5). OLLC Harmony reduction was used for trajectory inference ^149^. Trajectory inference was performed with Slingshot157 (v2.12.0), using the OLLC clusters as the input partition.

The start cluster was manually specified as OLLC_4, which showed a more progenitor-like marker profile and state (described in detail elsewhere in the manuscript), so that inferred lineages reflected progression away from an early OLLC precursor population, consistent with previous studies^114^. Slingshot-derived lineage-specific pseudotime values were computed for each cell, and an overall pseudotime measure (minimum pseudotime across lineages) was written back to the Seurat object for visualization and genotype-wise comparisons. For visualization of gene expression along OLLC pseudotime lineages, we used slingshot-derived pseudotime and curve weights to select cells belonging to the euploid and trisomic cluster–ending lineages. For selected oligodendrocyte-lineage genes, expression was plotted along pseudotime using GAM smoothing and projected onto the OLLC UMAP, with cells colored by normalized expression and lineage curves overlaid as dashed (euploid-ending) or solid (trisomic-ending) paths.

### Fate-conflict style pseudotime plots

To examine how these functional categories changed along oligodendrocyte-lineage trajectories, we generated “fate-conflict” style plots using Slingshot pseudotime^113^. For each category, we computed an aggregate module score per cell (mean of the corresponding module scores), scaled these scores between 0 and 1, and plotted smoothed curves of the category scores across pseudotime for selected Slingshot lineages, providing a trajectory-level view of how different transcriptional programs evolve within the OLLC population.

### Overlay with external reference datasets Age-specific alignment

To test whether the OLLC transcriptional states we identified in the FAs are also present in an independent dataset, we analyzed early postnatal human prefrontal cortex data^114^. We pooled oligodendrocytes and OPCs (Harmony_subclass_v5 annotation) from the human postnatal dataset^114^, to match the broader OLLC population analyzed in our FAs.

To identify the most developmentally comparable postnatal samples, we compared pseudobulk expression profiles between the FA OLLCs and the postnatal reference^114^. Average gene expression was calculated for euploid and trisomic FA-derived OLLCs and human OLLCs at each available age ≤200 days (euploid: days 60, 91, 120 and 142; trisomic: days 65 and 186). Spearman correlations were calculated across genes detected in both datasets with non-zero variance (n = 2000 genes), and each OLLC genotype was compared only with the matching postnatal genotype (euploid with control, trisomic with Ts21). Because each postnatal age represents a single sample, these correlations were used to assess relative transcriptional similarity rather than tested statistically.

Based on this analysis and the availability of closely age-matched samples from both genotypes (the reference human data set did not contain age matched controls for day 90 data point), we selected euploid cells from postnatal day 60 (554 OPCs and 207 oligodendrocytes; 761 cells) and trisomic cells from postnatal day 65 (643 OPCs and 66 oligodendrocytes; 709 cells) for subsequent analysis. These human cells were then projected onto the six FA-derived OLLC subclusters using Celda-derived gene modules to assess transcriptional correspondence between datasets.

### Module-based reference mapping

For each OLLC subcluster, we defined a transcriptional signature using its most enriched Celda modules, prioritizing co-regulated gene programs over individual markers or whole-transcriptome averages. This approach reduces sensitivity to single-cell data sparsity and broadly expressed genes, while rank- and specificity-based weighting improves discrimination between closely related OLLC states. Modules with at least five genes detected in the postnatal dataset were scored using Seurat AddModuleScore, which calculates expression relative to expression-matched control genes (https://search.r-project.org/CRAN/refmans/Seurat/html/AddModuleScore.html). We then calculated a weighted similarity score between each postnatal cell against each OLLC subcluster. Modules were assigned rank-based weights ranging from 5 (most enriched) to 1 (fifth most enriched). To limit the influence of programs shared across clusters, each weight was divided by the number of OLLC signatures that included that module. The weighted module scores were then normalized by the sum of their weights, yielding six similarity scores per cell.

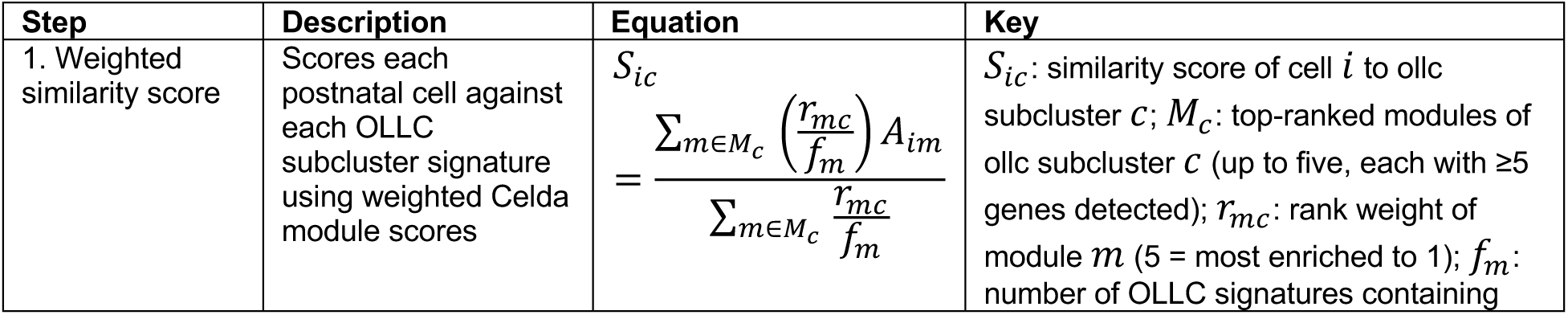
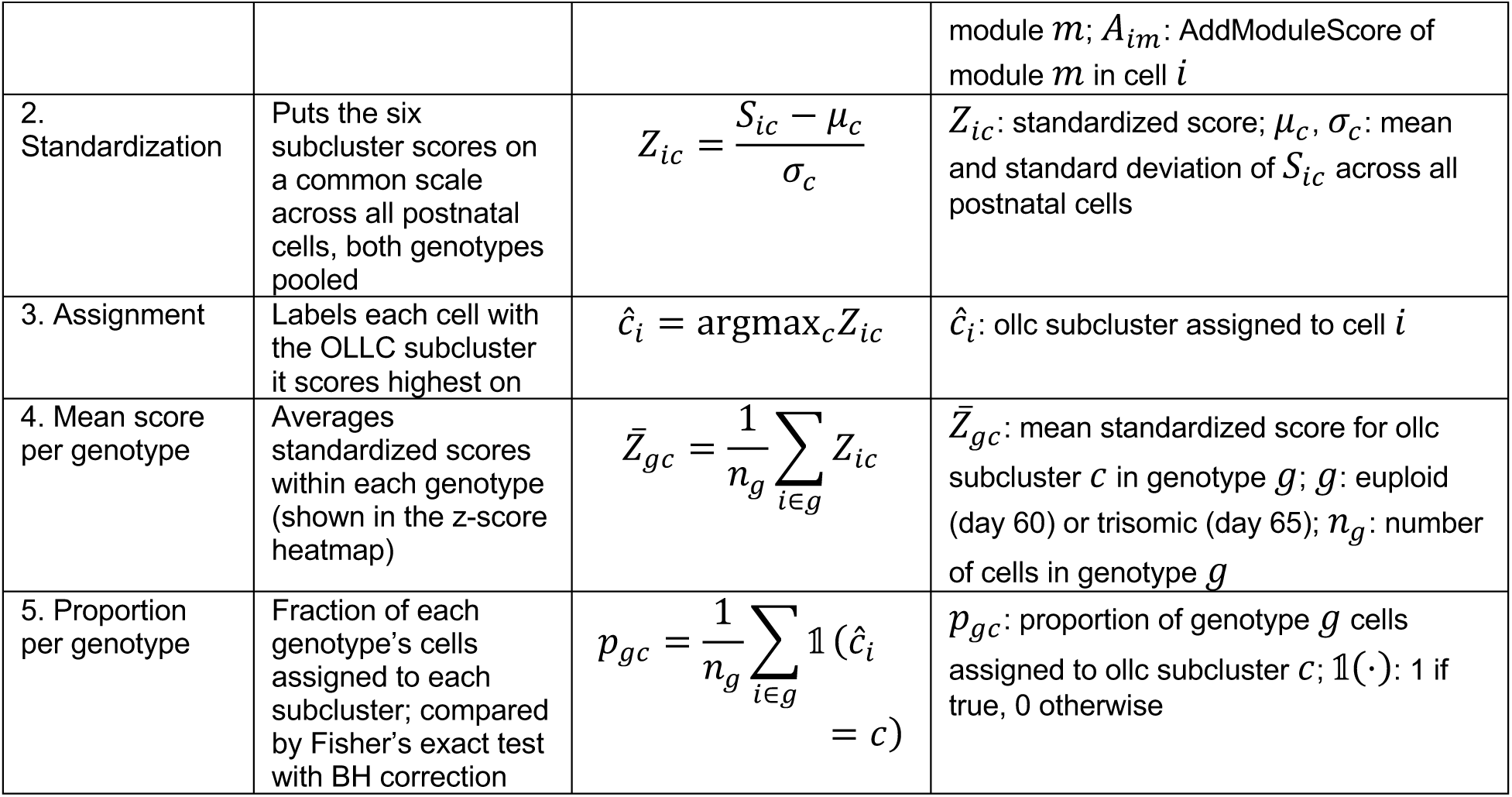

| Step | Description | Equation | Key |
|---|---|---|---|
| 1. Weighted similarity score | Scores each postnatal cell against each OLLC subcluster signature using weighted Celda module scores | $S_{ic} = \frac{\sum_{m\in M_c} \left(\frac{r_{mc}}{f_m}\right) A_{im}}{\sum_{m\in M_c} \frac{r_{mc}}{f_m}}$ | *S<sub>ic</sub>*: similarity score of cell *i* to ollc subcluster *c*; *M<sub>c</sub>*: top-ranked modules of ollc subcluster *c* (up to five, each with ≥5 genes detected); *r<sub>mc</sub>*: rank weight of module *m* (5 = most enriched to 1); *f<sub>m</sub>*: number of OLLC signatures containing module *m*; *A<sub>im</sub>*: AddModuleScore of module *m* in cell *i* |
| 2. Standardization | Puts the six subcluster scores on a common scale across all postnatal cells, both genotypes pooled | $Z_{ic} = \frac{S_{ic} - \mu_c}{\sigma_c}$ | *Z<sub>ic</sub>*: standardized score; *μ<sub>c</sub>*, *σ<sub>c</sub>*: mean and standard deviation of *S<sub>ic</sub>* across all postnatal cells |
| 3. Assignment | Labels each cell with the OLLC subcluster it scores highest on | *ĉ<sub>i</sub>* = argmax<sub>*c*</sub>*Z<sub>ic</sub>* | *ĉ<sub>i</sub>*: ollc subcluster assigned to cell *i* |
| 4. Mean score per genotype | Averages standardized scores within each genotype (shown in the z-score heatmap) | $\bar{Z}_{gc} = \frac{1}{n_g}\sum_{i\in g} Z_{ic}$ | *Z̄<sub>gc</sub>*: mean standardized score for ollc subcluster *c* in genotype *g*; *g*: euploid (day 60) or trisomic (day 65); *n<sub>g</sub>*: number of cells in genotype *g* |
| 5. Proportion per genotype | Fraction of each genotype's cells assigned to each subcluster; compared by Fisher's exact test with BH correction | $p_{gc} = \frac{1}{n_g}\sum_{i\in g} \mathbb{1}(\hat{c}_i = c)$ | *p<sub>gc</sub>*: proportion of genotype *g* cells assigned to ollc subcluster *c*; 𝟙(·): 1 if true, 0 otherwise |

Because raw scores were distributed differently for each OLLC signature, we standardized the scores for each subcluster across all postnatal cells, with both genotypes pooled. Each cell was then assigned to the OLLC subcluster with its highest standardized score, therefore reflecting relative enrichment for OLLC-derived transcriptional programs, rather than absolute cell identity or developmental-age equivalence between postnatal tissue and FAs. To compare signature enrichment between genotypes, we averaged the standardized similarity scores across all cells within each genotype, generating one mean score per OLLC subcluster for euploid (day 60) and trisomic (day 65) cells. Because scores were standardized across both genotypes pooled, positive values indicate higher-than-average enrichment for that OLLC signature in a given genotype, and negative values indicate lower-than-average enrichment.

We calculated the proportion of cells assigned to each OLLC subcluster within each genotype, so that proportions sum to one per genotype, and tested genotype differences per subcluster using Fisher’s exact test (assigned vs. not assigned × trisomic vs. euploid) with Benjamini–Hochberg correction across the six subclusters. Subclusters with FDR < 0.05 were considered enriched in the genotype with the higher proportion. These predicted distributions were compared with the genotype composition of the corresponding OLLC subclusters and with the independently inferred OLLC pseudotime trajectory. Because the module rankings and weights were derived from the OLLC dataset, this analysis represents a projection of FA-derived programs onto postnatal cells rather than an independent discovery analysis. Of note, genotype in the postnatal dataset is confounded with age (day 60 vs. day 65) and donor.

### CellChat Analysis

We used CellChat with the human ligand-receptor database, CellChatDB.human, to infer potential cell-cell communication between neuronal and oligodendroglial populations^165^. The analysis included ExNs (clusters 0, 1, 3, 6, 8, 9, 11, 13, 15, 17), InNs (clusters 18, 19, 20), and OLLCs. OLLC labels were updated using the Harmony-derived subclusters. For example, cells in Harmony cluster 0 were labeled OLLC_0, cells in Harmony cluster 1 were labeled OLLC_1, and the same approach was used for all OLLC clusters through OLLC_5. Neuronal cells were grouped into four broad categories: Immature ExNs (clusters 8, 9, 17), Mature ExNs (clusters 0, 1, 3, 6, 11, 13, 15), Immature InNs (cluster 18), and Mature InNs (clusters 19, 20).

CellChat was run separately for each sample using log-normalized RNA expression values. For each sample, the analysis identified expressed ligand-receptor pairs, estimated communication probabilities between cell groups, and summarized these interactions at the signaling-pathway level. We used the full human CellChat database, including secreted, extracellular matrix, and cell-contact signaling interactions. Communication probabilities were calculated with population.size= FALSE, so the inferred communication strength reflected ligand-receptor expression within each cell group rather than differences in cell-group abundance. Cell populations represented by fewer than three cells within a sample were excluded from the analysis. CellChat uses ligand and receptor expression to estimate potential communication between cell populations and organizes these interactions into broader signaling pathways^165^.

To compare genotype-associated signaling patterns, euploid and trisomic samples were evaluated as the same six matched cell line and sequencing-technology pairs used during the integration analysis. Within each pair, we compared pathway-level communication involving the euploid-enriched OLLC_2-TMS state in euploid samples and the trisomic-enriched OLLC_3-SID state in trisomic samples. Because each FA is either euploid or trisomic, OLLCs and neurons within a sample share the same genotype. Communication involving OLLC_2-TMS was therefore assessed with euploid neurons and communication involving OLLC_3-SID with trisomic neurons. This design preserves each cellular environment intact and captures differences in neuron–OLLC communication that may arise from either interacting partner. We examined signaling in both directions: neuronal populations to OLLCs and OLLCs to neuronal populations.

For each neuronal population, signaling pathway, and communication direction, we calculated the paired log_2_ fold change in inferred pathway probability for trisomic OLLCs relative to euploid OLLCs. Pathways were retained for visualization when the difference was in the same direction in at least three of the available matched pairs. For these pathways, chord plots display the median inferred pathway probability across matched pairs for OLLC_2 and OLLC_3, with edge width representing the median pathway probability. Because CellChat is based on ligand and receptor gene expression, these results should be interpreted as predicted communication potential rather than direct evidence of physical cell-cell interactions or functional signaling.

### CellChat Discrepancy Plots

To further examine communication involving genotype-enriched OLLC states, CellChat interaction counts and aggregate communication strengths were extracted for neuronal population to OLLC signaling and OLLC to neuronal population signaling. In euploid samples, analyses focused on OLLC_2, whereas in trisomic samples, analyses focused on OLLC_3. For each sample and neuronal population, we extracted the number of retained ligand-receptor interactions and the total inferred communication strength for both signaling directions.

For interactions showing the same direction of difference in at least half of the matched euploid and trisomic comparisons, median values were calculated across the matched pairs separately for OLLC_2 and OLLC_3. These median interaction counts and communication strengths were used for the discrepancy plots and chord plots. The median was used to represent the typical value across paired comparisons while limiting the influence of individual samples with unusually high or low inferred communication values.

### CellChat consensus pathway heatmaps

To identify reproducible differences in inferred CellChat signaling, pathway-level communication probabilities were compared within matched euploid and trisomic sample pairs. Analyses were performed separately for signaling from neurons to genotype-enriched OLLC states, and for signaling from genotype-enriched OLLC states to neurons. For each neuronal sender or receiver population and each CellChat signaling pathway, paired log_2_ fold changes were calculated as the inferred pathway probability in OLLC_3 relative to OLLC_2. A pseudocount of 1 × 10^6^ was added before log transformation to allow calculation of fold changes when a pathway probability was zero in one sample. Differences were summarized across available matched comparisons using the median paired log_2_ fold change.

CellChat infers communication probabilities from the average expression of ligands and receptors within each cell group rather than from the number of cells, and population size was not used to weight probabilities (population.size = FALSE). Differences in the number of cells per OLLC state or neuronal population therefore do not directly drive differences in inferred probability, and downsampling was not performed. In addition, all comparisons were made within matched euploid–trisomic sample pairs and summarized across pairs rather than on pooled data, which limits the influence of differences in sample size between individual samples.

Pathways were selected based on reproducibility at the pathway level rather than for individual ligand–receptor (L-R) pairs. For each signaling direction and neuronal population, a pathway was considered consistent if the direction of difference agreed in at least three matched comparisons (the circle-plot criterion). Comparisons with zero inferred probability were included, and only pairs lacking the relevant cell populations were excluded. From these consistent pathways, biologically relevant pathways were further assessed at the L-R level and shown as bubble plots. Within each of these pathways, sender–receiver combinations consistent in at least three matched comparisons are displayed, together with L-R pairs detected in at least one sample, using values from the genotype-associated OLLC state with higher pathway-level signaling. L-R values were averaged across samples rather than summarized as medians, so that interactions detected in only a subset of samples were not reduced to zero.

### CellChat Circle Plots

Consensus CellChat circle plots were generated to visualize reproducible signaling from genotype-enriched OLLC states to neuronal receiver populations. For each CellChat signaling pathway, inferred pathway probabilities were extracted from the matched euploid and trisomic sample comparisons. Euploid samples contributed communication from OLLC_2-TMS, while trisomic samples contributed communication from OLLC_3-SID.

For each neuronal receiver population and signaling pathway, euploid and trisomic pathway probabilities were summarized across matched pairs using the median of the raw CellChat probability values. Median values were used to provide a robust summary of the typical inferred communication strength across pairs while reducing the influence of an individual pair with an unusually high or low value. Only neuronal receiver and pathway combinations showing the same direction of difference in at least three matched comparisons were included.

A separate circle plot was generated for each retained signaling pathway. In each plot, OLLC_2-TMS and OLLC_3-SID were shown as sender nodes and neuronal populations were shown as receiver nodes. Edges from OLLC_2-TMS represent the median inferred pathway probability in euploid samples, and edges from OLLC_3-SID represent the median inferred pathway probability in trisomic samples. Edge width corresponds to the median pathway probability across matched pairs. All neuronal populations were retained as nodes in each plot, including populations without a consensus-supported interaction for a given pathway, to allow direct comparison of network structure across signaling pathways.

Only pathways with at least one consensus-supported OLLC-to-neuron interaction were visualized. Companion tables reported the number of evaluable matched pairs, the number supporting higher signaling in each genotype-associated OLLC state, median pathway probabilities, interquartile ranges, and the median paired log_2_ fold change for each displayed edge.

### Ligand receptor plots

To identify the ligand-receptor (L-R) pairs underlying consensus-supported pathways, we generated CellChat-style bubble plots for selected signaling pathways in both directions of communication: from OLLCs to neuronal populations and from neuronal populations to OLLCs. As for the circle plots, euploid samples contributed communication involving OLLC_2-TMS and trisomic samples contributed communication involving OLLC_3-SID, and neuronal populations included ExN, immature ExN, InN, and immature inhibitory InN.

### Cellchat Alignment with Reference Dataset

To test whether the genotype-associated differences in OLLC–neuron communication we observed in FAs also occur in human tissue, we ran CellChat on the same early postnatal reference cells used for module-based mapping (euploid postnatal day 60, trisomic postnatal day 65)^114^. Postnatal OPCs and oligodendrocytes were labeled by their assigned OLLC subcluster, and we grouped all excitatory and inhibitory neuronal subclasses into single ExN and InN categories (Harmony_subclass_v5 annotation). CellChat was run separately for each genotype using the same settings as for the FA data. As in the FA analysis, communication involving cells assigned to OLLC_2-TMS was assessed in euploid samples, and communication involving cells assigned to OLLC_3-SID in trisomic samples. For each signaling pathway, direction (neuron to OLLC or OLLC to neuron) and neuronal population, we asked whether pathway-level communication was stronger for the OLLC_3-SID or OLLC_2-TMS assigned state in the postnatal data, and compared this with the consensus direction from our FA matched-pair analysis. Because the postnatal dataset does not separate immature neurons, we restricted this comparison to mature ExNs and InNs. Each FA consensus call was then classified as concordant (same direction in the postnatal data), discordant (opposite direction), or not detected.

## Acknowledgments

The authors thank Dr. Yuriy Alekseyev and the members of the Boston University Chobanian & Avedisian School of Medicine Microarray and Sequencing Resource Core for their help and guidance with the sequencing experiments. We appreciate Madeline Kuppe-Fish from UMass Chan for snRNA-seq sample processing guidance and we thank Shiraz Bheda from Neural Stem Cell Institute for invaluable input on the transcriptomic analysis. We are also thankful to Dr. Andre Sousa, from the University of Wisconsin–Madison, and Drs. Celine Vuong and Luis de la Torre-Ubieta, from the University of California, Los Angeles, for sharing data used in their original publications^114,115^ and helpful discussions. We acknowledge the technical assistance of Lorenz Ponce in EM tissue processing, block sectioning, and grid preparation and give special thanks to Aarnav Kandukuri, Musarat Khan, Vianne Lim, and Gloria Olubowale for manual cell tracking. We are grateful to NIH agencies for their support: NIH/NIA RF1AG088529 (MPI: E. Zeldich and M. Thunemann), NIH/NINDS R21NS125469 (MPIs: E Zeldich and M. Medalla), NIH/NIA and R03NS126864 (PI: E. Zeldich). Research reported in this publication was supported by the National Science Foundation Award CBET 2215990. The content is solely the responsibility of the authors and does not necessarily represent the official views of the National Science Foundation.

## Declaration of generative AI and AI-assisted technologies in the writing process

During the preparation of this work the authors used in ChatGPT (OpenAI) and Claude (Anthropic) to assist with language editing, proofreading, and improving clarity of the manuscript. After using this tool, the authors reviewed and edited the content as needed and take full responsibility for the content of the published article.

## Author contributions

NBC designed and performed all aspects of the cell culture experiments and interpreted the results. NBC, SK, and EZ wrote the first drafts of the manuscript. EZ and MM conceived the idea and oversaw the project, provided guidance, interpreted the results, and edited the manuscript. SK and JDC designed and performed bioinformatic analyses for scRNA-seq and snRNA-seq datasets. NBC, GF, SC, CC, ML, and MK performed image acquisition and quantitative analysis of immunostaining. NBC performed live-cell imaging for manual cell tracking, MT, MW, and CH developed analysis pipeline, and NXC analyzed live-cell imaging data. NBC and MM performed electron microscopy tissue processing and imaging, and NBC and MK performed electron microscopy image analysis.

## Resource availability

Raw and processed scRNA-seq and snRNA-seq data generated in this study have been deposited in the NCBI Gene Expression Omnibus (GEO) under accession number GSE352389. The human fetal cortex reference data are available from GEO under accession number GSE305146. Early postnatal human prefrontal cortex data were obtained from Risgaard et al. 2026^114^.

## Code availability

All code used for data processing and analysis is available at https://github.com/Sonaldk2311/FA_Trisomy21_OLLC and archived at Zenodo (https://doi.org/10.5281/zenodo.23776722).

