## Supplementary Figure for "Trisomy 21 Perturbs Oligodendrocyte Developmental States and Oligo–Neuron Communication in Human Forebrain Assembloids"

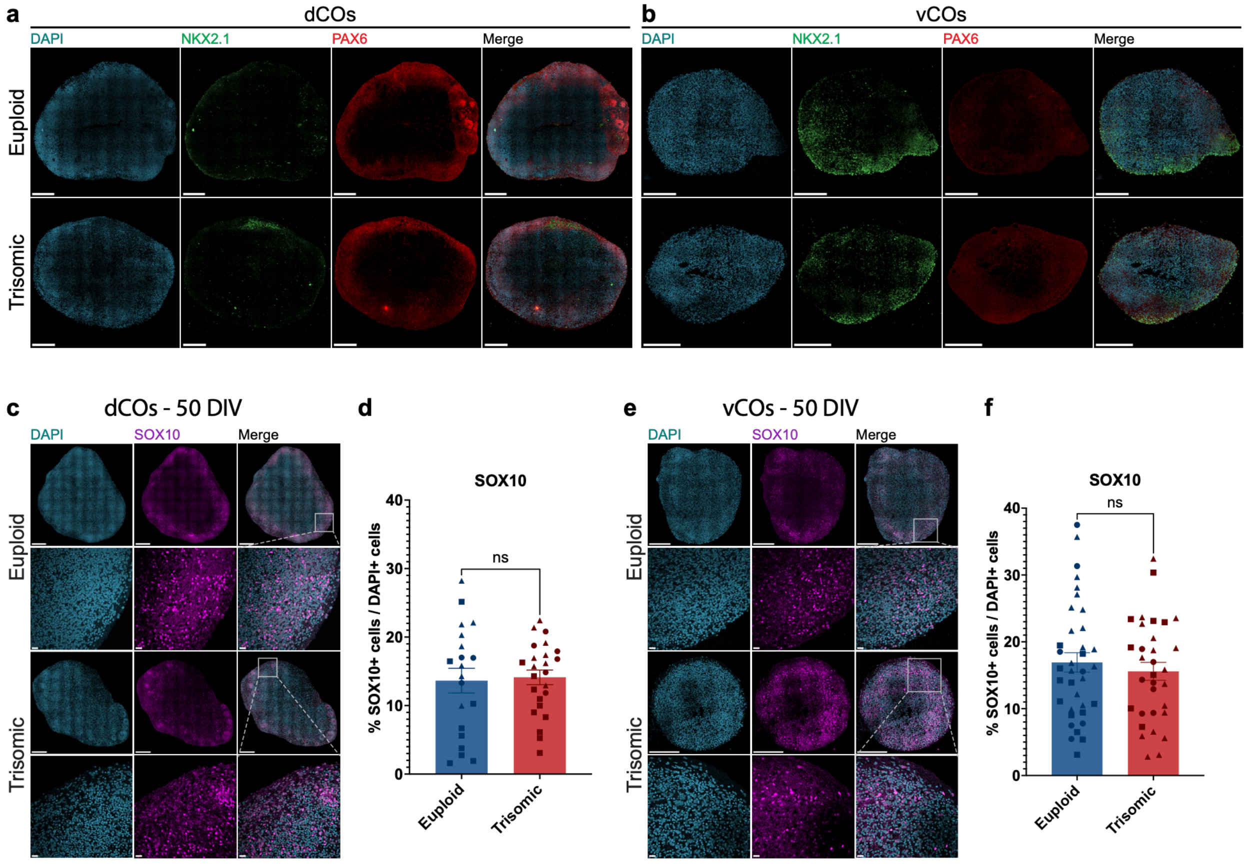


*Supplementary Figure 1: Regional identity and oligodendrocyte lineage specification in euploid and trisomic cortical organoids (COs) before fusion.*

**a, b,** Representative high-resolution stitched maximum intensity projection confocal images showing regional identity markers in euploid and trisomic (**a**) dorsal COs (dCOs) and (**b**) ventral COs (vCOs) at 50 days *in vitro* (50 *DIV*). Dorsal identity is indicated by the high expression of PAX6 (red) and low expression of the ventral lineage marker NKX2.1 (green). Ventral identity is indicated by the high expression of NKX2.1 and low expression of PAX6, confirming regional patterning at 50 *DIV*. Nuclei are counterstained with DAPI (blue). Scale bar, 300 µm. **c–f,** Representative stitched and inset confocal images from euploid and trisomic dCOs (**c**) and vCOs (**e**) at 50 *DIV* showing SOX10 (magenta) expression. Scale bars, 300 µm (overviews) and 20µm (insets). **d, f,** Quantification of percentage of SOX10+ cells in (**d**) dCOs (LME, $\beta = +0.3802$, $p = 0.8029$) and (**f**) vCOs (LME, $\beta = -1.332$, $p = 0.4990$) within total DAPI+ cells per CO. Euploid, $n = 20$. Trisomic, $n = 24$ dCOs; Euploid, $n = 36$. Trisomic, $n = 32$ vCOs. Graphs display results obtained from two independent differentiation experiments. All comparisons were analyzed using linear mixed-effects models, with genotype effect reported in the figure legend and the effects of cell line and differentiation summarized in **Supplementary Table 1**. Each point represents one dCO or vCO and each symbol shape represents an isogenic cell line pair. circle = WC; square = ILD; triangle = DS1. Results shown as mean ± SEM. ns = not significant.

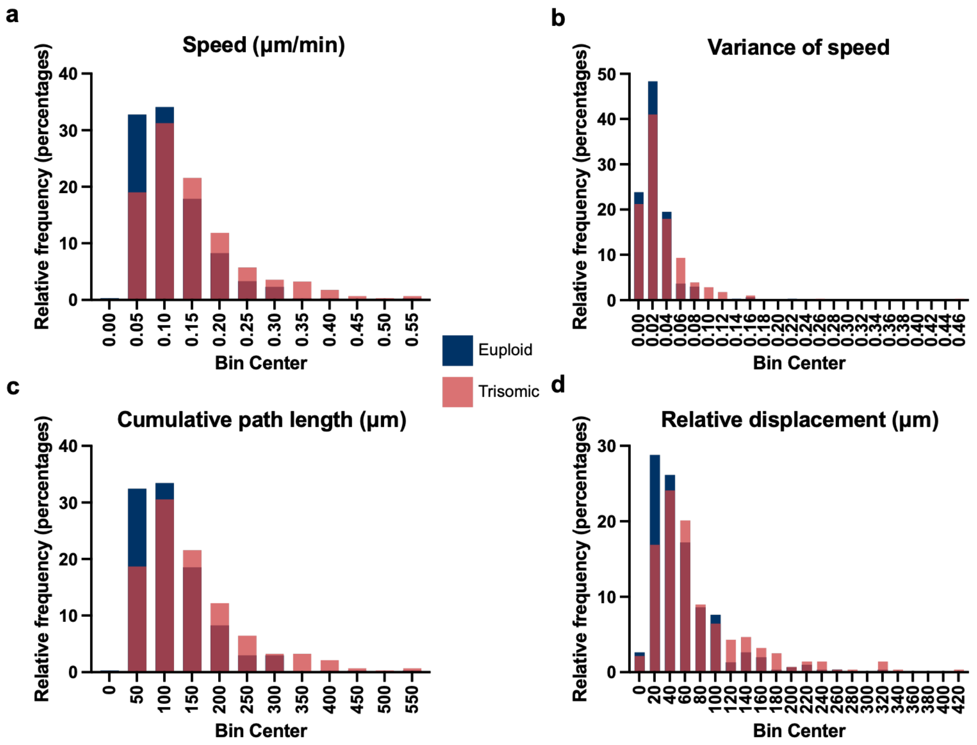


*Supplementary Figure 2: Distribution of migration measurements in euploid and trisomic SOX10-TdTomato+ OLLCs in FAs at 120 DIV.*

**a–d**, Frequency distributions of measurements obtained from individually tracked *SOX10-tdTomato*+ cells in euploid and trisomic FAs. Histograms show (**a**) cell migration speed (μm/min), (**b**) variance of speed, (**c**) cumulative path length (μm), and (**d**) relative displacement (μm) values. Values were binned and expressed as relative frequencies (percentages) of tracked cells within each genotype. Euploid and trisomic values are overlaid to show differences in distributions. Blue = euploid; red = trisomic. Euploid, *n* = 302. Trisomic, *n* = 278 *SOX10-TdTomato*+ cells.

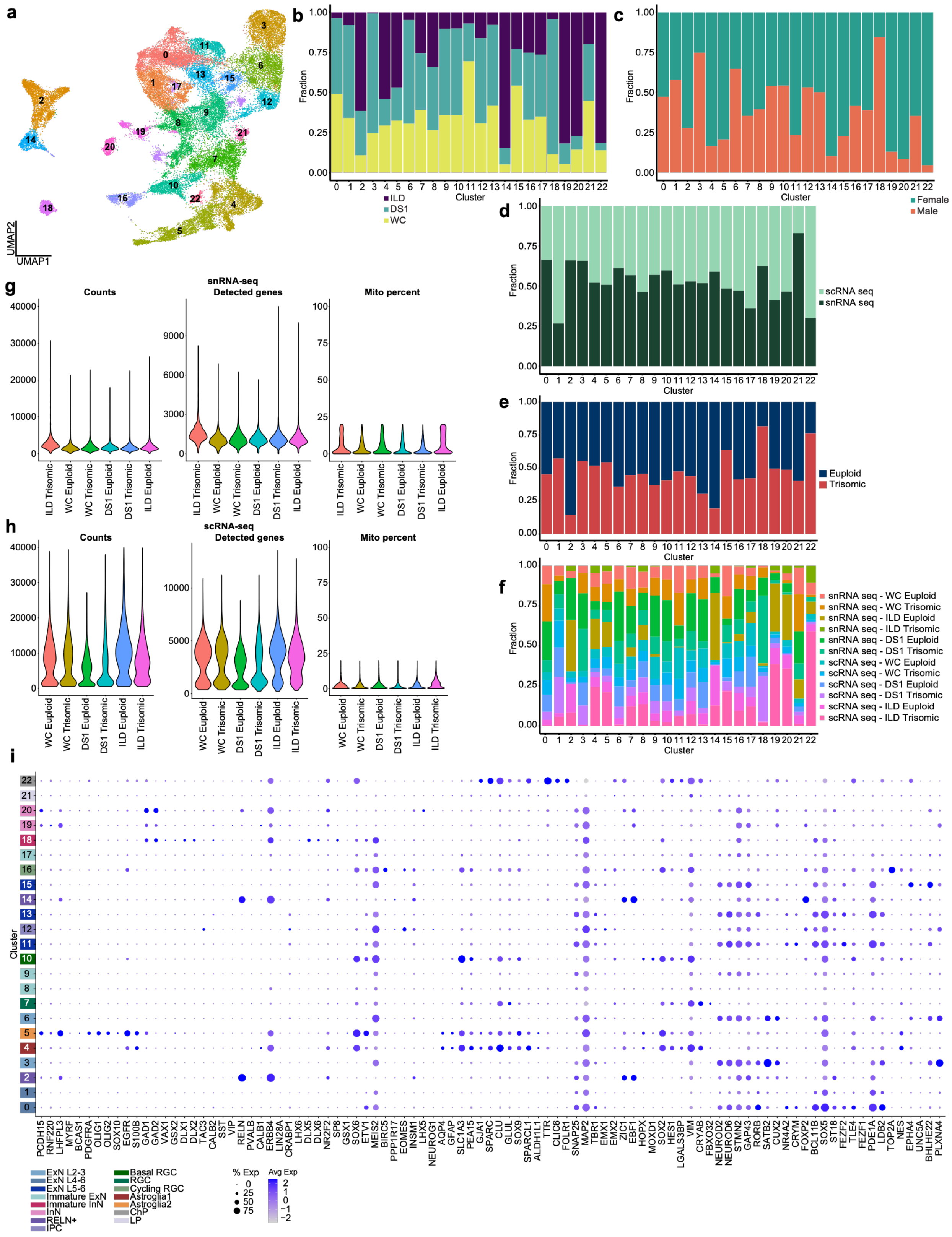


*Supplementary Figure 3: Integration, quality control, and cell-type annotation of single-cell and single-nucleus transcriptomic datasets obtained from FAs at 150 DIV.*

**a,** Uniform manifold approximation and projection (UMAP) of the integrated snRNA-seq and scRNA-seq dataset (45,547 cells and nuclei) from all cell lines, genotypes and sequencing technologies. Colors and numbers indicate the 23 clusters (0–22) identified by unsupervised graph-based clustering of the Harmony-integrated object. **b–f**, Stacked bar plots showing the fraction of cells in each cluster derived from each (**b**) cell line (WC, ILD, DS1), (**c**) sex, (**d**) sequencing technology (scRNA-seq, snRNA-seq), (**e**) genotype (euploid, trisomic) and (**f**) sample (sequencing technology × cell line × genotype). **g, h,** Violin plots of quality control metrics per sample for (**g**) snRNA-seq nuclei and (**h**) scRNA-seq cells, showing the number of unique molecular identifier (UMI) counts, the number of detected genes, and the percentage of mitochondrial transcripts per cell. **i**, Dot plot of marker genes used to annotate cell types across clusters. Clusters (rows) are labeled and color-coded by assigned cell type (legend, bottom left). Dot size indicates the percentage of cells in each cluster expressing the gene, and color indicates average scaled expression.

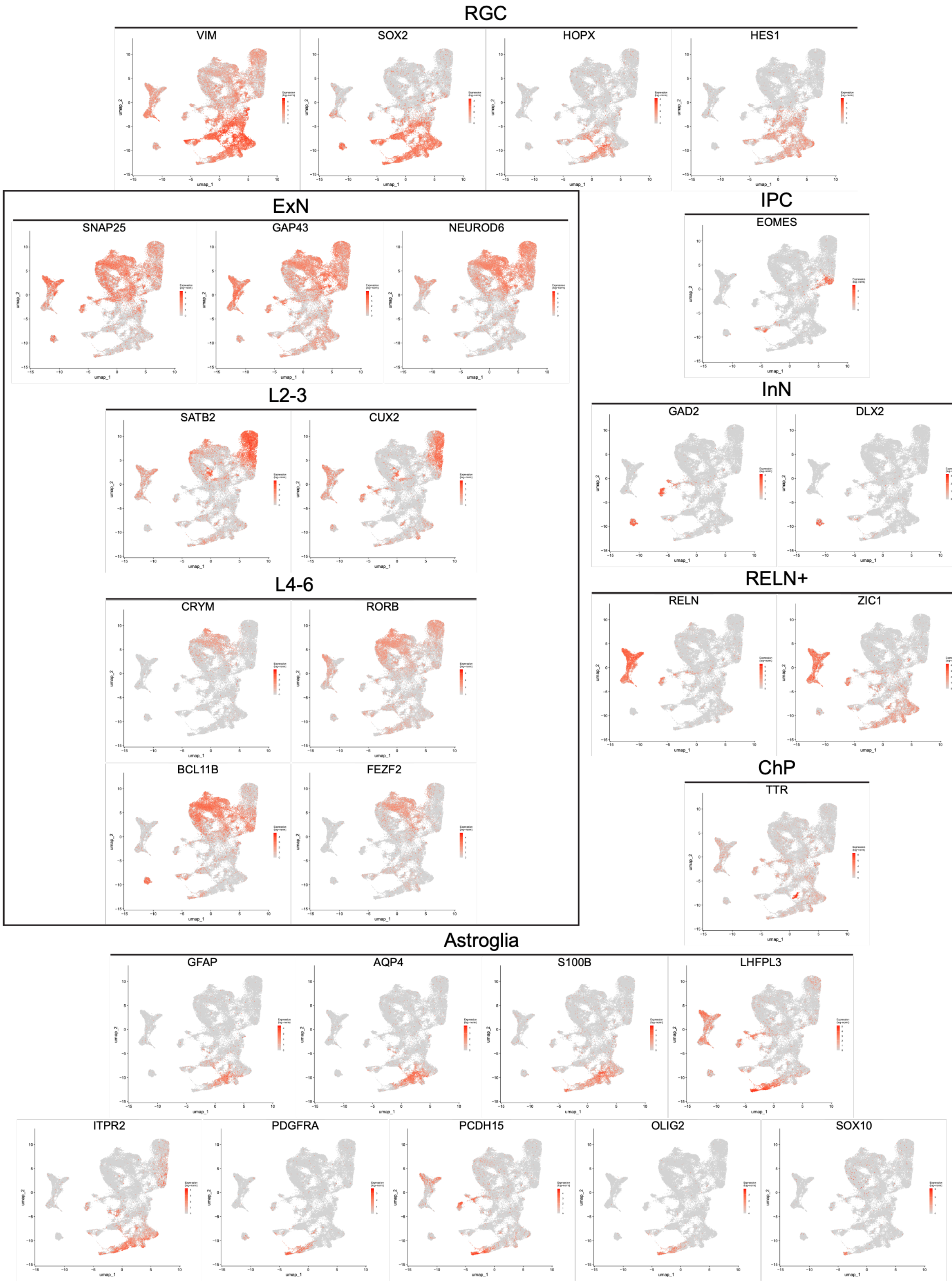


*Supplementary Figure 4: Expression of canonical cell-type markers across the integrated transcriptomic dataset.*

UMAP feature plots showing expression of canonical markers used for cluster annotation in the integrated scRNA-seq and snRNA-seq dataset based on markers in **Supplementary Table 2.** Expression levels are displayed using the color scale indicated in each plot, with gray representing cells with low or undetected expression.

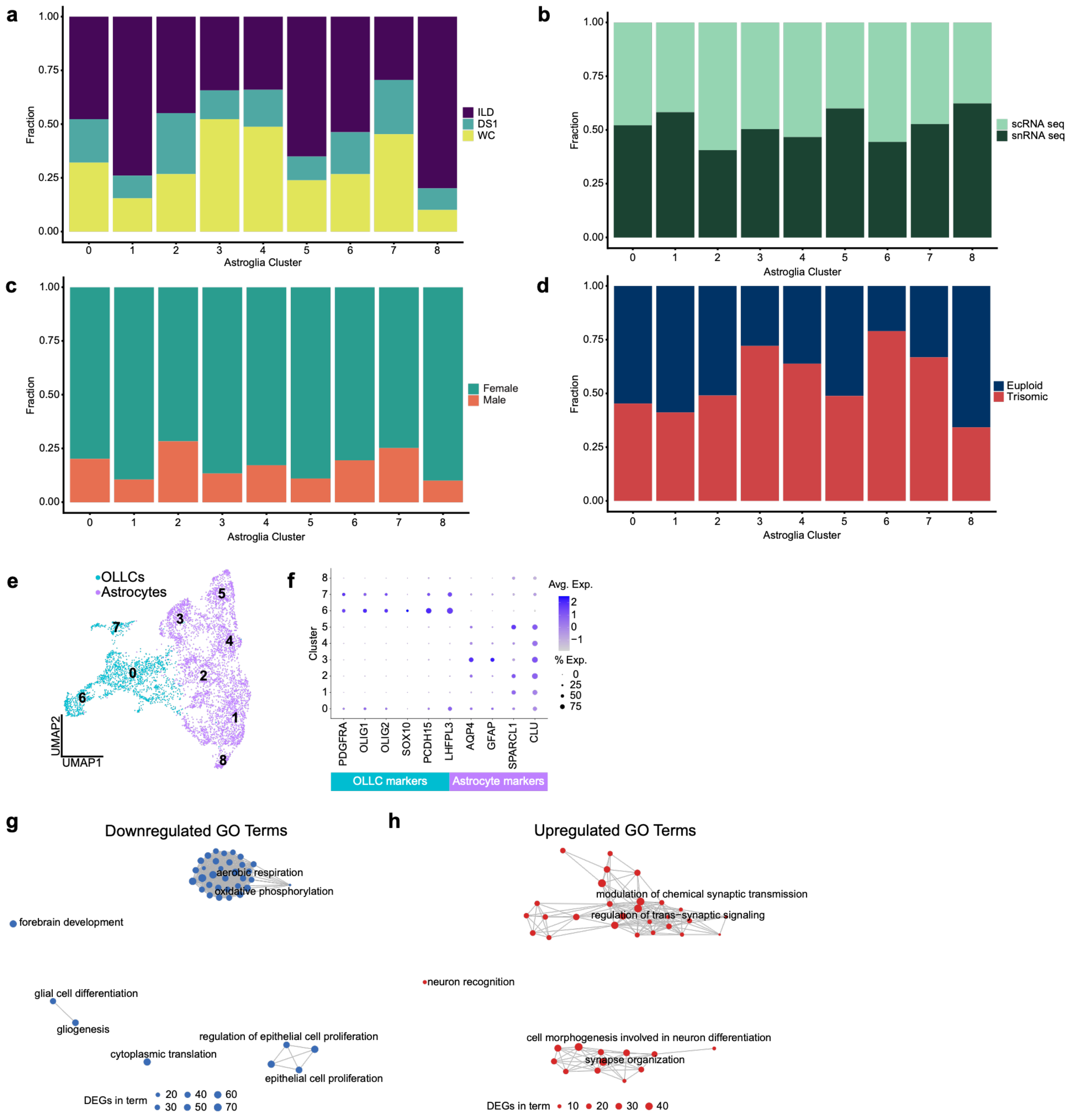


*Supplementary Figure 5: Astroglial subclustering segregates OLLCs in FAs.*

**a–d,** Stacked bar plots showing the fraction of (**a**) cell line, (**b**) sequencing technology, (**c**) sex, and (**d**) genotype in each astroglial subcluster. **e,** UMAP representation of astroglial subset from the Astroglia1 and Astroglia2 clusters of the integrated transcriptomic dataset and reclustered to resolve heterogeneity within the astroglial compartment. Subclusters assigned to OLLCs and astrocytes are indicated by distinct colors. **f**, Dot plot showing native Seurat visualization of canonical astroglial and OLLC markers across astroglial subclusters (0–8). Subclusters 0, 6, and 7 were annotated as OLLCs based on *PDGFRA*, *OLIG1*, *OLIG2*, *SOX10, PCDH15,* and *LHFPL3* expression, whereas subclusters 1, 2, 3, 4, 5, and 8 were annotated as astrocytes based on expression of *AQP4, GFAP, SPARCL1*, and *CLU*. Dot size corresponds to the percentage of cells expressing each gene in a cluster, and color indicates average scaled expression. **g, h,** Network plots of GO Biological Process terms downregulated (**g**) and upregulated (**h**) in trisomic vs. euploid OLLCs. DEGs were identified with scMetaIntegrator across six matched cell line × sequencing technology pairs (Fisher FDR < 0.05). The 40 most significant terms (BH-adjusted p < 0.05) are shown. Node size indicates the number of DEGs per term, and edges connect terms that share genes.

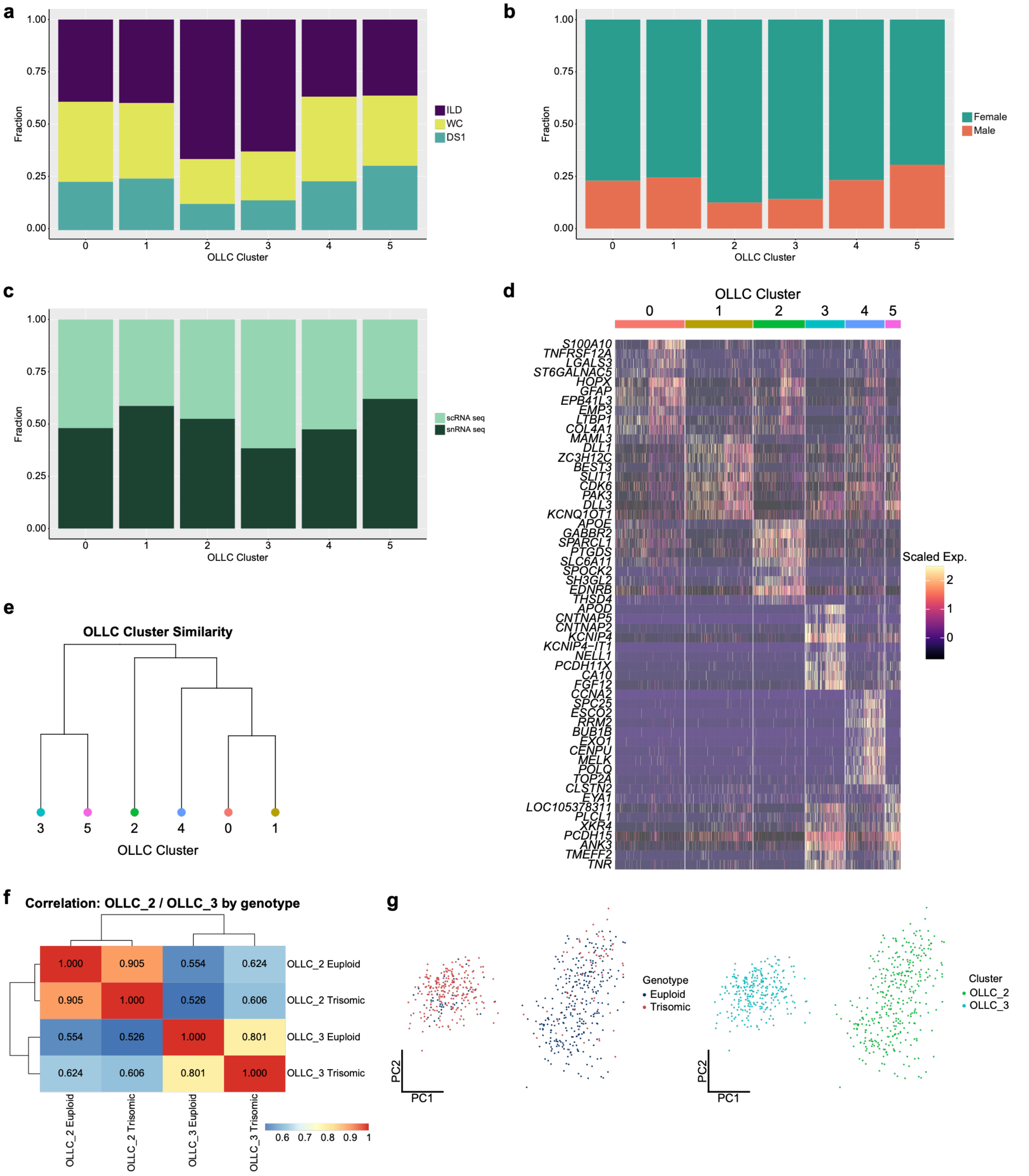


*Supplementary Figure 6: Transcriptional characterization and similarity of OLLC subclusters.*

**a–c,** Stacked bar plots showing the fraction of (**a**) cell line, (**b**) sex, and (**c**) sequencing technology in each of the six OLLC clusters (OLLC_0–OLLC_5). **d**, Heatmap of the top marker genes for each OLLC subcluster. Markers were identified with Seurat FindAllMarkers (Wilcoxon rank-sum test, each cluster versus all other OLLCs), restricted to the 1,155 highly variable genes shared across all cell line × sequencing technology groups, and filtered for FDR < 0.05, average $\log_2$ fold change > 0.25, and expression in < 50% of cells outside the corresponding subcluster. Up to 10 genes with the highest average $\log_2$ fold change are shown per cluster. Columns are individual cells grouped by subcluster; color indicates scaled expression. **e,** Hierarchical clustering dendrogram of OLLC subclusters based on the mean Harmony embedding (dimensions 1–15) of each cluster, using Euclidean distance and complete linkage (Seurat BuildClusterTree). **f,** Heatmap showing pairwise Spearman correlations between pseudobulk transcriptional profiles of euploid and trisomic cells within OLLC_2-TMS and OLLC_3-SID. Correlation coefficients are indicated within each tile, and hierarchical clustering illustrates transcriptional similarity between the four populations. **g,** Principal component analysis (PCA) of cells assigned to OLLC_2-TMS and OLLC_3-SID, colored by genotype (left; euploid, blue; trisomic, red) or OLLC state (right; OLLC_2-TMS, cyan; OLLC_3-SID, green). Each point represents an individual cell.

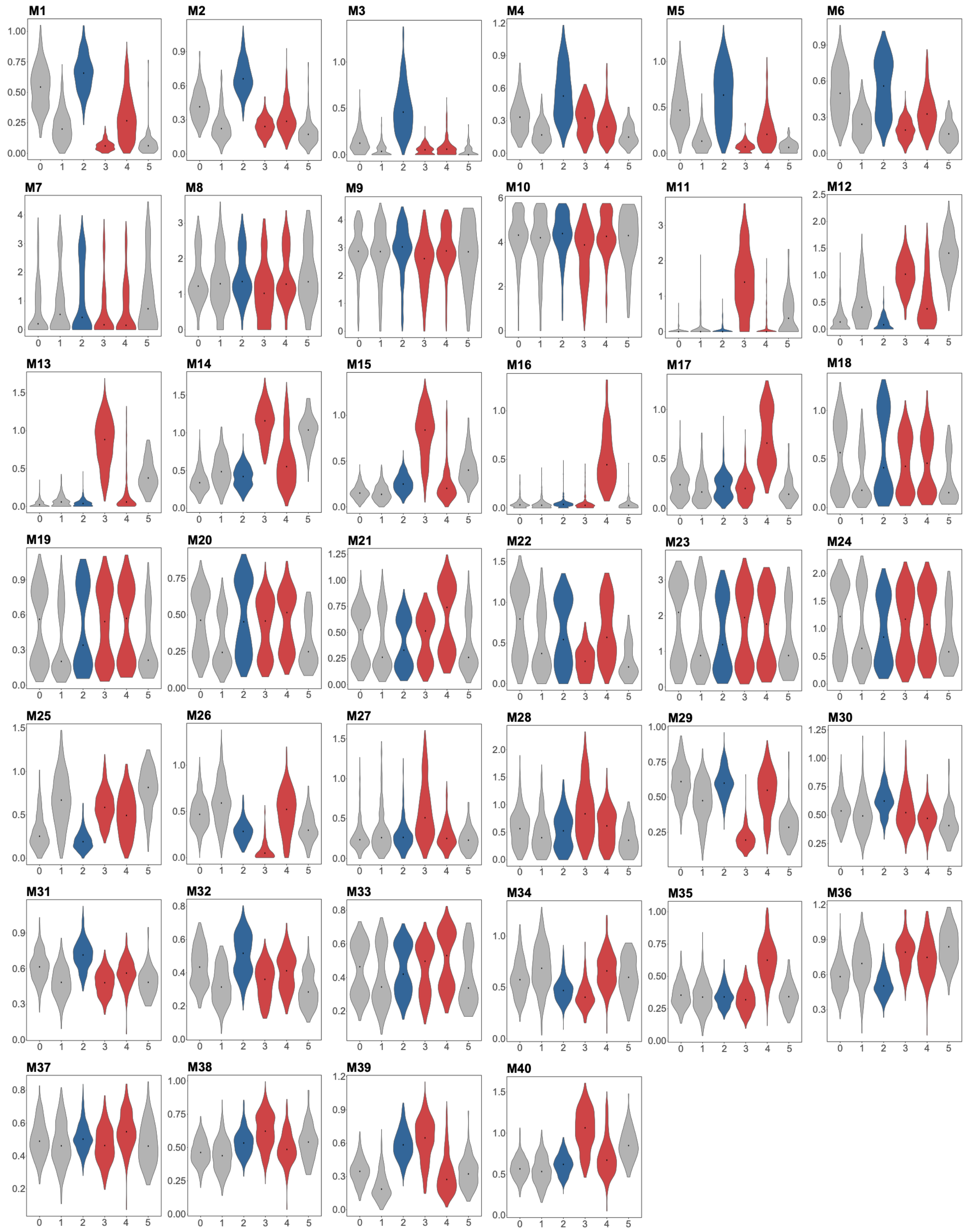


*Supplementary Figure 7: Distribution of Celda gene module expression across OLLC subclusters.*

Violin plots for each Celda gene module (M1–M40) showing the distribution of per-cell module expression across OLLC subclusters, calculated as the mean log-normalized expression of all module genes in each cell. Dots indicate the median. Subclusters are colored by genotype enrichment: blue, euploid-enriched (OLLC_2); red, trisomic-enriched (OLLC_3 and OLLC_4); gray, not enriched (OLLC_0, OLLC_1 and OLLC_5).

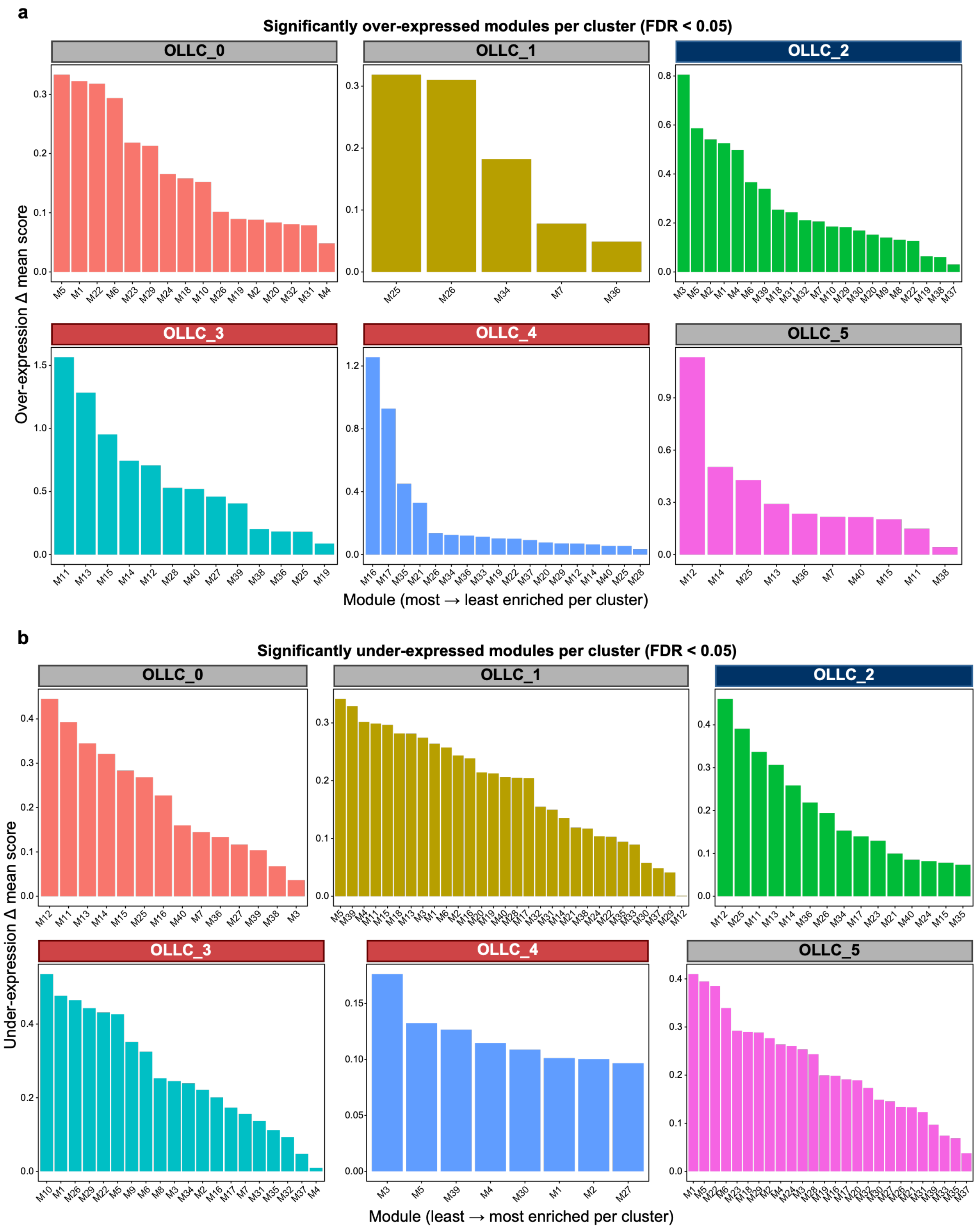


*Supplementary Figure 8: Differential enrichment and depletion of Celda gene modules across OLLC subclusters.*

**a, b,** Bar plots showing (**a**) significantly overexpressed and (**b**) underexpressed Celda gene modules in each OLLC subcluster (OLLC_0–OLLC_5). For each cell, a module score was calculated as the mean z-scored expression of the genes in that module. Module scores for each cluster were compared with those of all other OLLCs using Wilcoxon rank-sum tests, and p values were adjusted with the Benjamini–Hochberg method. All modules with FDR < 0.05 are shown, with a (**a**) positive or (**b**) negative difference in mean module score. Significantly overexpressed modules were defined as those with FDR < 0.05 and a positive mean difference in module score, and significantly underexpressed modules as those with FDR < 0.05 and a negative mean difference in module score. Bar height indicates the magnitude of the difference in mean module score between the cluster and all other OLLCs, and modules are ordered from largest to smallest difference within each cluster. Subclusters are colored by genotype enrichment: blue, euploid-enriched; red, trisomic-enriched; gray, not enriched.

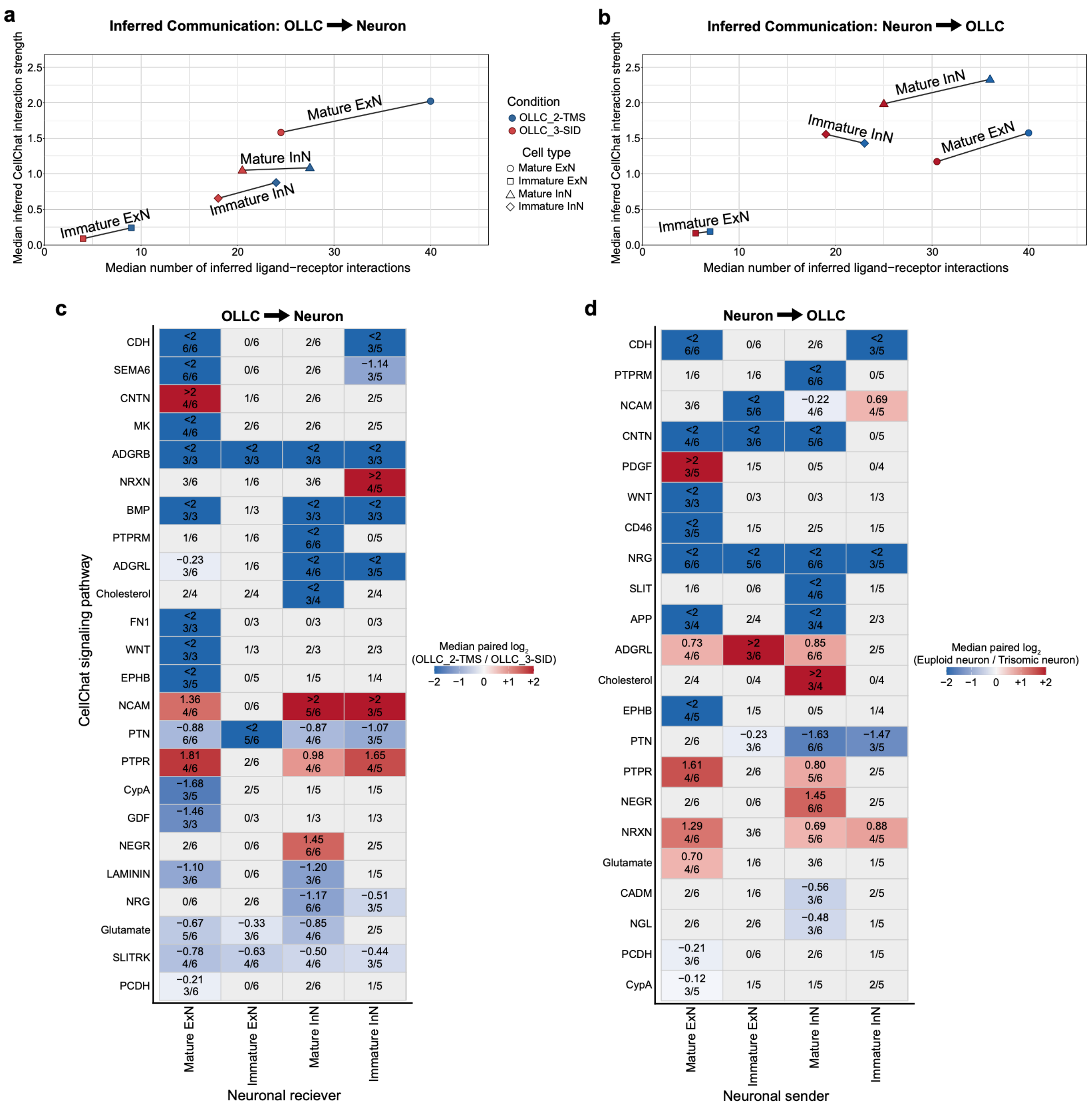


**Supplementary Figure 9:** OLLC-state-dependent differences in inferred neuron–OLLC communication.

**a, b,** Comparative CellChat discrepancy plots showing inferred communication between OLLC states and neuronal populations, analyzed separately for (**a**) OLLC-to-neuron and (**b**) neuron-to-OLLC signaling. Each point represents interactions involving one neuronal population (mature or immature excitatory neurons (ExNs) or inhibitory neurons (InNs)). The x-axis shows the median number of retained ligand-receptor interactions and the y-axis shows the median aggregate inferred communication strength across matched pairs. Blue represents communication involving

OLLC_2-TMS in euploid samples, and red represents communication involving OLLC_3-SID in trisomic samples. Point shapes distinguish neuronal populations, and connecting lines link corresponding neuronal populations across the two conditions. Circle, Mature ExN; square, Immature ExN; triangle, Mature InN; diamond, Immature InN. **c, d,** Heatmaps showing differences in inferred CellChat signaling pathways for (**c**) OLLC-to-neuron communication, with neuronal populations as receivers, and (**d**) neuron-to-OLLC communication, with neuronal populations as senders. Communication involving OLLC_2-TMS was inferred in euploid samples and communication involving OLLC_3-SID in trisomic samples, each with neurons from the same samples. For each pathway and neuronal population (mature and immature ExNs and InNs), paired $\log_2$ fold changes in inferred pathway probability (OLLC_3-SID relative to OLLC_2-TMS) were calculated within each of the six matched cell line × sequencing-technology pairs. Tile color shows the median paired $\log_2$ fold change (red, higher inferred signaling involving OLLC_3-SID; blue, higher inferred signaling involving OLLC_2-TMS). The color scale is capped at ±2, and values beyond this range are labeled "< -2" or "> +2". Text within each tile represents the median $\log_2$ fold change and the number of matched pairs supporting that direction out of the number of evaluable pairs. Colored tiles indicate pathway-population combinations with a consistent direction in at least three matched pairs; the grey tiles indicate combinations that did not meet this criterion. Pathways were ordered according to the largest absolute median effect observed across neuronal populations.

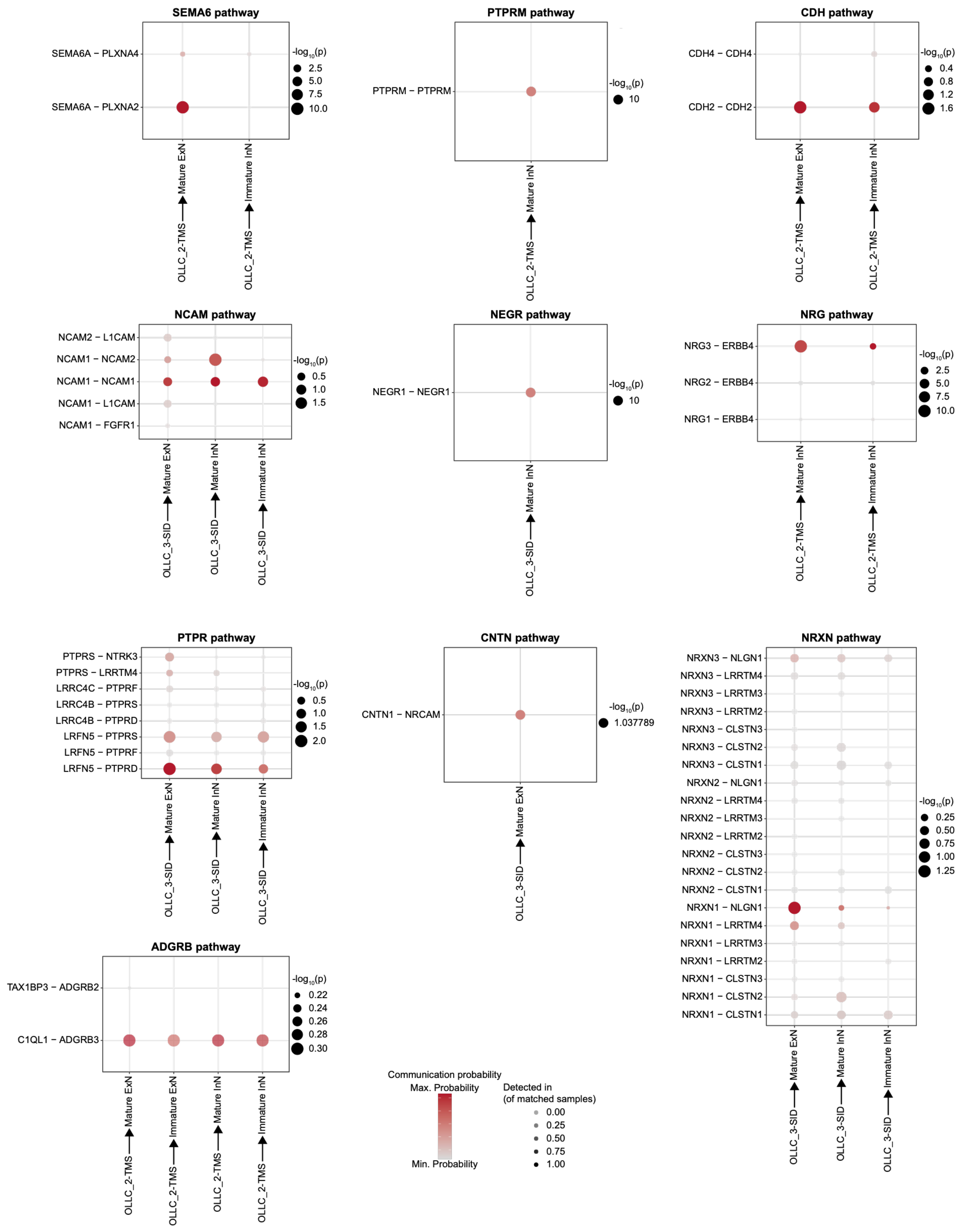


*Supplementary Fig. 10: Representative individual ligand-receptor (L-R) interactions underlying OLLC-to-neuron signaling pathways.*

Bubble plots showing inferred L-R interactions within selected CellChat signaling pathways involved in communication from OLLC_2-TMS or OLLC_3-SID to neuronal populations. Pathways include SEMA6, PTPRM, CDH, NCAM, NEGR, NRG, PTPR, CNTN, NRXN, and ADGRB. Each plot displays individual L–R pairs (rows) and the neuronal receiver populations in which the corresponding pathway met the consensus criterion of a consistent direction of difference in at least three matched cell line × sequencing technology comparisons (columns). Interactions are shown for OLLC_2-TMS in euploid samples or OLLC_3-SID in trisomic samples, according to the OLLC state with higher inferred pathway-level signaling. Bubble color represents the mean inferred CellChat communication probability across samples, bubble size indicates $-\log_{10}(p)$ for the inferred L–R interaction, and shading indicates the proportion of matched samples in which the interaction was detected. These plots illustrate the individual L-R interactions contributing to the pathway-level differences summarized in **Supplementary Fig. 9c**.

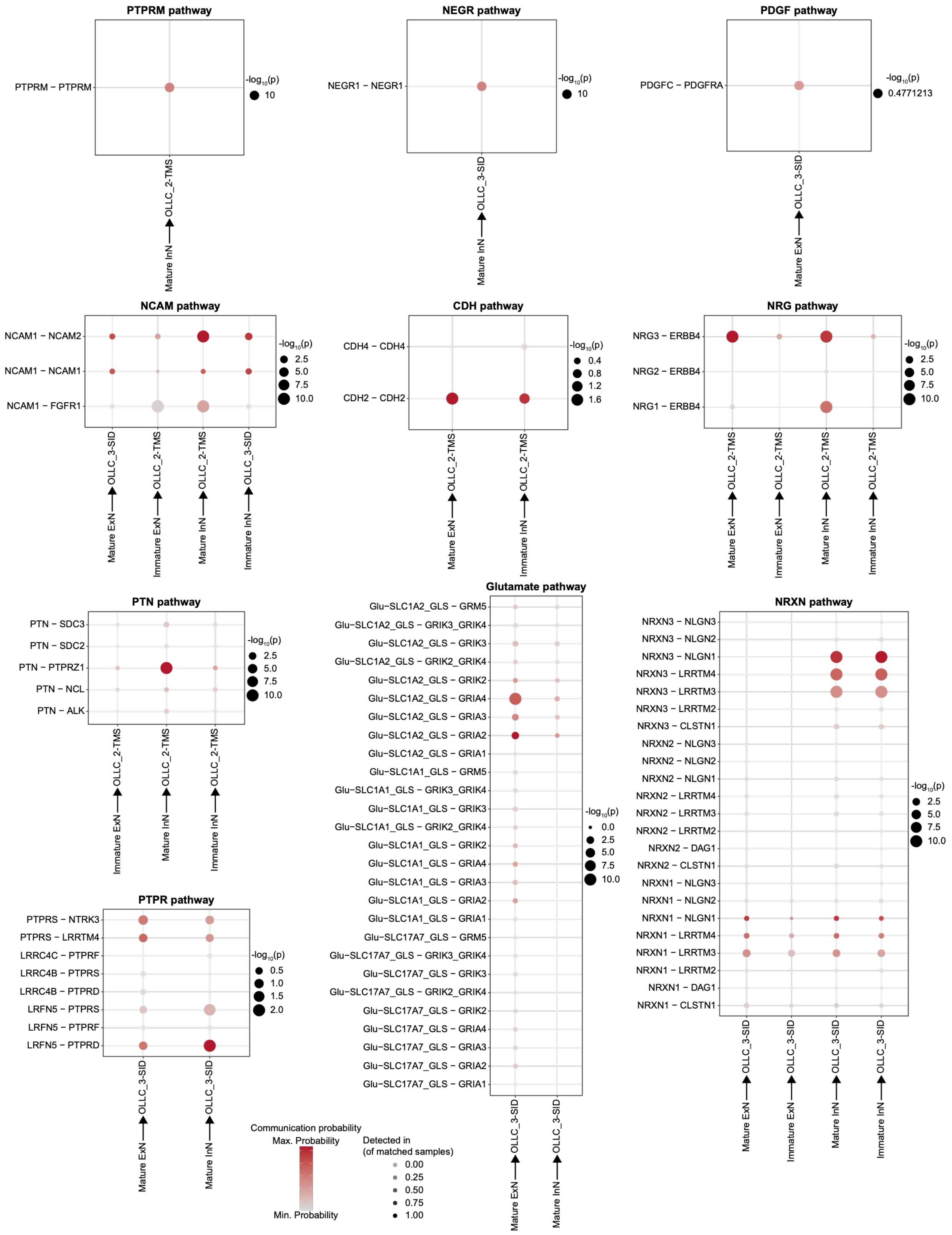


*Supplementary Fig. 11: Representative individual ligand-receptor (L-R) interactions underlying neuron-to-OLLC signaling pathways.*

Bubble plots showing inferred L–R interactions within selected CellChat signaling pathways involved in communication from neuronal populations to OLLC_2-TMS or OLLC_3-SID. Pathways include PTPRM, NEGR, PDGF, NCAM, CDH, NRG, PTN, PTPR, Glutamate, and NRXN. Each plot displays individual L-R pairs (rows) and the neuronal sender populations in which the corresponding pathway met the consensus criterion of a consistent direction of difference in at least three matched cell line × sequencing technology comparisons (columns). Interactions are shown for neuronal populations from euploid samples communicating with OLLC_2-TMS or from trisomic samples communicating with OLLC_3-SID, according to the condition with higher inferred pathway-level signaling. Bubble color represents the mean inferred CellChat communication probability across samples, bubble size indicates $-\log_{10}(p)$ for the inferred L–R interaction, and shading indicates the proportion of matched samples in which the interaction was detected. These plots illustrate the individual L-R interactions contributing to the pathway-level differences summarized in **Supplementary Fig. 9d**.

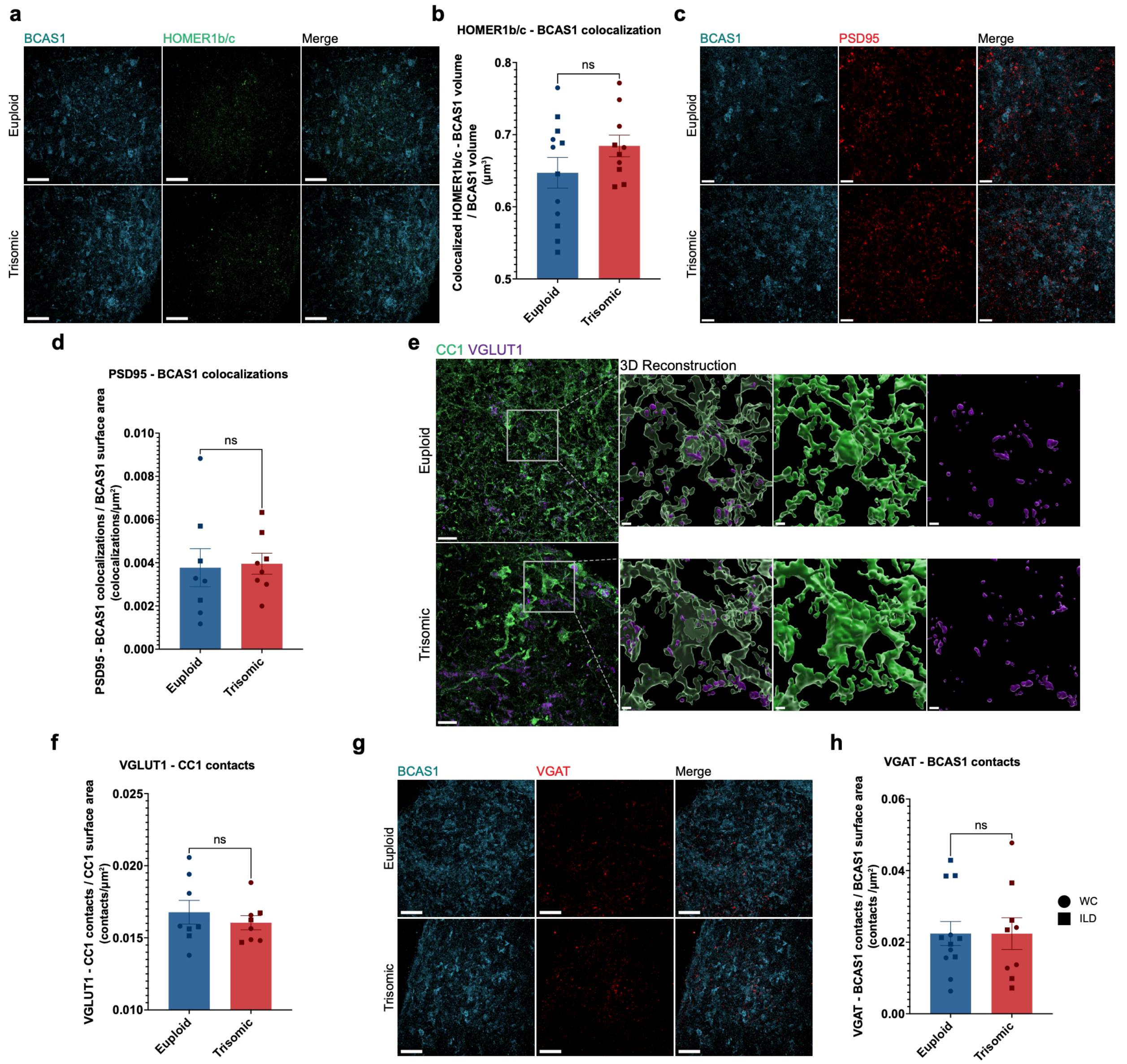


*Supplementary Figure 12: Synaptic marker associations with oligodendrocyte lineage cells in euploid and trisomic FAs at 150 DIV.*

**a,** High resolution confocal images from euploid and trisomic FAs showing BCAS1 (blue) and HOMER1b/c (green). Scale bar, 50 µm. **b,** Quantification of colocalized HOMER1b/c-BCAS1 volume (defined as overlapping expression of both proteins > 0.01µm$^3$) normalized to BCAS1 volume (LME; β = +0.0351, *p* = 0.1448) per FA. Euploid, *n* = 12. Trisomic, *n* = 10 FAs. **c,** High resolution confocal images showing BCAS1 (blue) and PSD95 (red). Scale bar, 20 µm. **d,** Quantification of PSD95-BCAS1 colocalizations (defined as PSD95 puncta within 0.01 µm of BCAS1 expression) normalized to BCAS1 surface area (colocalizations/µm$^2$; LME; β = +0.0004, *p* = 0.6037) per FA. Euploid, *n* = 8. Trisomic, *n* = 8 FAs. **e,** High resolution confocal images and corresponding reconstructions of CC1 and VGLUT1 expression in trisomic and euploid FAs. Scale bars, 50 µm (confocal) and 3 µm (reconstruction insets) **f,** Quantification of VGLUT1-CC1 contacts (defined as VGLUT1 expression within 1 µm of CC1 expression) normalized to CC1 surface area (contacts/µm$^2$; LME; β = -0.0004*,* *p* = 0.5653) per FA. Euploid, *n* = 8. Trisomic, *n* = 8 FAs. **g,** Representative confocal images from euploid and trisomic FAs showing BCAS1 (blue) and VGAT (red) Scale bar, 50 µm. **h,** Quantification of VGAT-BCAS1 contacts (defined as VGAT puncta within 1 µm of BCAS1 expression) normalized to BCAS1 surface area (contacts/µm$^2$; LME; β = +0.0009, *p* = 0.8535) per FA. Euploid, *n* = 12. Trisomic, *n* = 9 FAs. Graphs display results from two independent differentiation experiments. All comparisons were analyzed using linear mixed-effects models, with genotype effect reported in the figure legend and the effects of cell line and differentiation summarized in **Supplementary Table 1**. Each point represents one FA and each symbol shape represents an isogenic cell line pair. circle = WC; square = ILD Results shown as mean ± SEM. ns = not significant.

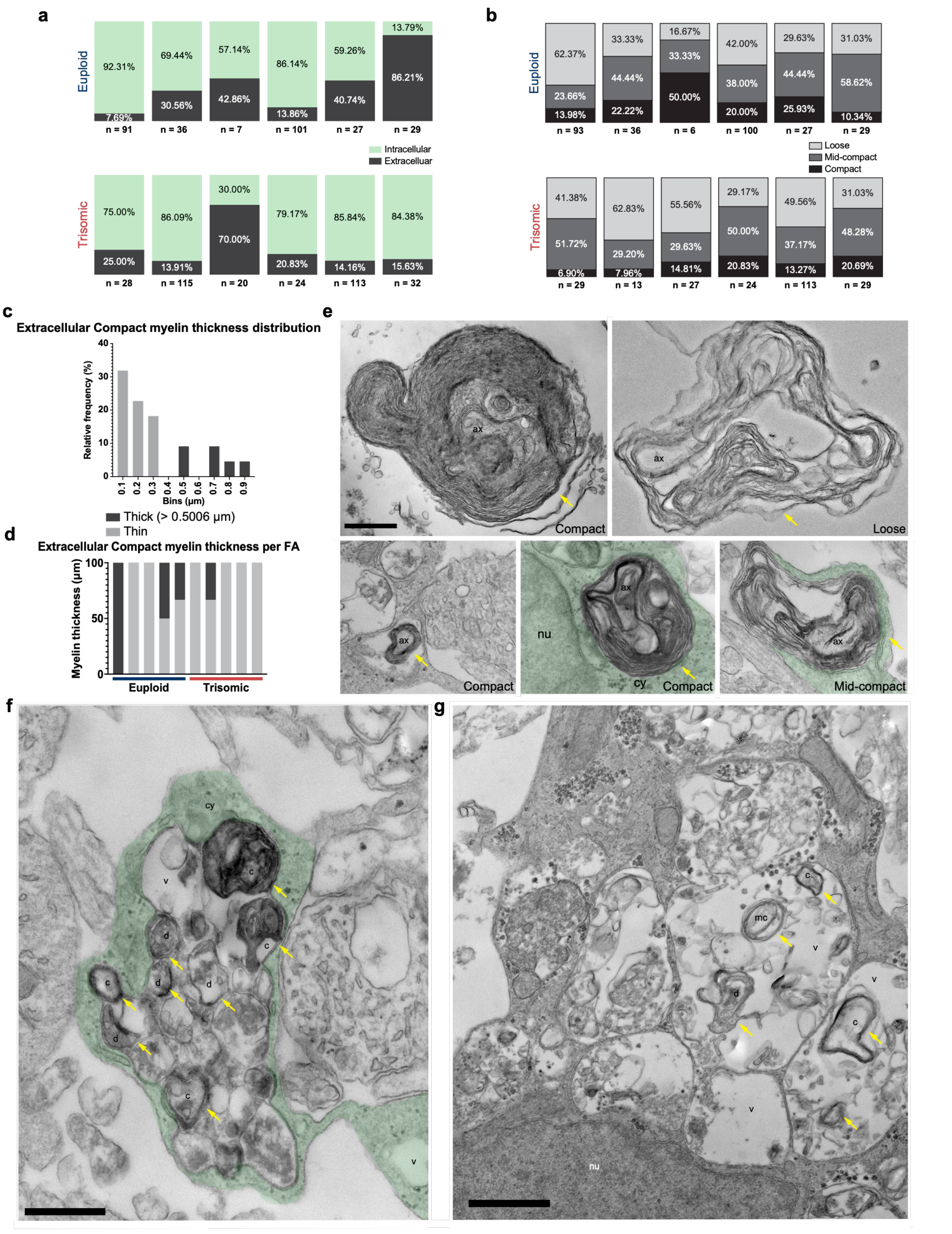


*Supplementary Figure 13: Ultrastructural analysis of myelin organization and compaction in FAs.*

**a, b,** Stacked bar graphs showing (**a**) intracellular (green) and extracellular (dark grey) myelin and (**b**) loose (light grey), mid-compact (dark grey) and compact (black) myelin proportions in individual FAs. **c**, Frequency distribution showing extracellular thick (dark grey) and thin (light grey) compact myelin proportions in all extracellular compact myelin. **d,** Stacked bar plot showing extracellular thick (dark grey) and thin (light grey) compact myelin proportions in individual euploid (blue) and trisomic (red) FAs. Thick myelin is defined as myelin thicker than 0.5006 μm. **e,** Electron micrographs showing loose and compact extracellular myelin in FAs at 150 *DIV,* and mid-compact and compact intracellular myelin, surrounded by cytoplasm (green). Scale bar, 500 nm. **f, g,** Electron micrographs showing defective intracellular and extracellular myelin. Scale bars, 500 nm (**f**) and 1 μm (**g**). ax = axon; cy = cytoplasm (green); nu = nucleus; v = vacuole; c = compact myelin; mc = mid-compact myelin; d = defective myelin. Yellow arrows indicate myelin.
